# Molecular mechanism of mitochondrial phosphatidate transfer by Ups1/Mdm35

**DOI:** 10.1101/702415

**Authors:** Jiuwei Lu, Kevin Chan, Leiye Yu, Jun Fan, Yujia Zhai, Fei Sun

**Author notes:** These authors make equal contribution to this work (Jiuwei Lu, Kevin Chan, Leiye Yu). Correspondence (F.S.), (Y.Z.) or (J.F.).

## Abstract

Cardiolipin plays many important roles for mitochondrial physiological function and is synthesized from phosphatidic acid (PA) at inner mitochondrial membrane (IMM). PA synthesized from endoplasmic reticulum needs to transfer to IMM via outer mitochondrial membrane (OMM). The transfer of PA between IMM and OMM is mediated by Ups1/Mdm35 protein family. Although there are many structures of this family available, the detailed molecular mechanism of how PA is transferred between membranes is yet unknown. Here, we report another crystal structures of Ups1/Mdm35 in the authentic monomeric apo state and the DHPA bound state. By combining subsequent all-atom molecular dynamics simulations, extensive structural comparisons and biophysical assays, we discovered the conformational changes of Ups1/Mdm35, identified key structural elements and residues during membrane binding and PA entry. We found the monomeric Ups1 on membrane is an important transit for the success of PA transfer, and the equilibrium between monomeric Ups1 and Ups1/Mdm35 complex on membrane affects the PA transfer rate and can be regulated by many factors including environmental pH.

## INTRODUCTION

As the powerhouse of cell, mitochondria produce energy through oxidative phosphorylation, a process in which ATP is created via electron transfer chain and ATP synthase embedded in the inner membrane of mitochondria (IMM). IMM has a very high concentration of cardiolipin (CL) that accounts for about 20% of the total lipids of IMM ^1^. CL not only interacts with a number of IMM proteins, but also serves as an integral component of respiratory complexes and participates in their folding ^2^. In recent years, the role of CL in mitochondrial signaling pathways has also been highlighted. Upon stress signals, CL is externalized to the outer membrane of mitochondria (OMM) forming a binding platform for the specific recruitment of signaling molecules that are required for mitophagy and apoptosis ^3^. In eukaryotes, CL is synthesized on the matrix side of IMM via an enzymatic cascade starting from phosphatidic acid (PA) ^4^. PA is a rare component of the mitochondrial membrane (< 1% of the total lipids) that is predominantly synthesized in endoplasmic reticulum (ER) and imported to OMM ^1, 5^. Although it remains unclear how PA is transferred from ER to OMM, the hetero-dimeric protein complex, termed Ups1/Mdm35 in yeast or PRELID1/TRIAP1 in mammalian cells, has recently been identified as a lipid transfer complex that mediates the transport of PA from OMM to IMM ^6–10^.

Ups1 is located in mitochondrial intermembrane space (IMS) and was identified as a member of the conserved UPS1/PRELI family proteins, which are related to mitochondrial function ^11^. Ups1 was initially found to affect the biogenesis of Mgm1, which is the homologue of human OPA1 and required for mitochondrial fusion ^12^. Cells lacking Ups1 showed dramatically reduced amounts of short-form Mgm1 (s-Mgm1) and had less tubular mitochondria ^12^. Deletion of Ups1 resulted in a considerable decrease in CL level ^13^. However, this decrease could be restored by simultaneous deletion of Ups2, which is the homolog of Ups1 in yeast. Mdm35 is another conserved yeast IMS protein and belongs to the twin Cx9C protein family ^14^. As the binding partner of the Ups proteins, Mdm35 not only facilitates their import into mitochondria, but also protects them against proteolysis ^15, 16^. The role of Ups1/Mdm35 complex in lipid transfer was discovered in 2012 and for the first time Ups1 was recognized as a lipid transfer protein that can shuttle PA between mitochondrial membranes^8^. In addition, the dynamic assembly of Ups1 with Mdm35 was found to be essential for the PA transfer process. The closest homologue of Ups1/Mdm35 in human is PRELID1/TRIAP1 that facilitates PA transfer in vitro^9^. In addition to PRELID1, human possesses another three Ups1 homologues named PRELID2, SLMO1 and SLMO2, all of which shares the conserved “PRELID” domain with PRELID1 and form a complex with TRIAP1 (a homologue of Mdm35) ^10^.

In 2015, two groups reported crystal structures of Ups1/Mdm35 with or without PA, in which the interaction between Ups1 and Mdm35 and the PA binding pocket of Ups1 were revealed ^17, 18^. In the same year, Miliara et al reported the crystal structure of SLMO1/TRIAP1, which is structurally similar with Ups1/Mdm35. Despite no significant sequence homology, Ups1/Mdm35 as a whole shows a remote structural similarity to other lipid transfer proteins (LTPs) including the phosphatidylinositol transfer proteins (PITPs) and the related cholesterol-binding START domains ^19, 20^. Structural comparison highlights the α2-loop in Ups1 that seems to be the equivalent region of the lipid exchange loop of PITP-α or the Ω1-loop in the START domains ^17, 21^. These two loops were thought to be the membrane-docking site for lipid loading and unloading and function as a lid of the lipid-binding cavity. The outward and inward movements of the lid cause the widening and closing of the cavity entrance, respectively, which help the lipid enter/exit from the cavity or to be retained in the cavity ^19, 22^. In the apo and lipid-bound PITP-α structure, the position of this lid is dramatically different, producing a distinct open and closed state of the lipid-binding cavity. However, the situation seems different in Ups1/Mdm35 system. Although experiments showed the importance of the α2-loop for Ups1 function, in which deleting this loop or replacing all residues with Ala or mutating conserved L62 and W65 into Ala dramatically impaired the PA transfer activity of Ups1/Mdm35, it is surprising that the conformation of the α2-loop has little change between the apo and PA-bound Ups1/Mdm35 structures ^17, 18^. Furthermore, Ups1 mutant without the α2-loop could still bind to liposomes containing CL, suggesting the role of α2-loop might not be relevant to the membrane-docking site ^17^. Mutagenesis experiments revealed some conserved basic residues located either at the bottom of the PA binding pocket (R25 and R54), or at the periphery near the PA binding site (H33, K58, K61, K148 and K155) are critical for PA transfer^18, 21^. In addition, it was found that the dissociation of Mdm35 could enhance the binding of Ups1 to the membrane ^17^. Recently, crystal structures of human PRELID1-TRIAP1 and PRELID3b-TRIAP1 were solved and the structural determinants of lipid specificity were studied by yeast genetic screen and coarse-grain simulations^23^.

These studies extend our understanding of the PA transfer mechanism between mitochondrial membranes. However, they also raised more questions about the exquisite details of the PA transfer process. For example, how does Ups1/Mdm35 or Ups1 bind to the membrane? How does Mdm35 dissociate from Ups1? How does PA enter or exit from the PA binding pocket? What is the exact role of the α2-loop? Does it undergo a conformational change before and after PA binding? How is the PA transfer regulated? In the present study, we combined structural approach, molecular dynamics (MD) simulations and biophysical assays to further elucidate the molecular mechanism of PA transfer by Ups1/Mdm35. Based on our studies and previous reports, we also proposed a working molecular model for Ups1/Mdm35 mediated PA transport and performed the kinetic analysis.

## RESULTS

### Structure of monomeric Ups1/Mdm35 complex

The crystal structure of SeMet substituted *S. cerevisiae* Ups1/Mdm35 (PDB code 5JQL) was determined at a resolution of 2.9 Å by single-wavelength anomalous dispersion method (Supplementary Table 1). The asymmetric unit of the crystal contains six Ups1/Mdm35 heterodimers. The overall structures of heterodimers are similar except that the positions of the C-terminal long α3-helices of Ups1 orient slightly differently (Supplementary Figs. 1 and 2). The C-terminal long α3-helices (residues K138-E168) of every two Ups1 (Ups1A and Ups1B) exchange with each other to form a domain-swapped dimer (Fig.1a). This result is consistent with the previously reported structure (PDB code 4YTW) by Y. Watanabe et al ^17^. By gel-filtration analysis and cysteine crosslinking assay, they found the domain-swapped dimer likely arose from the crystallization artifact, as a result they built an artificial monomer for the subsequent structural analysis ^17^. However, it’s unclear whether this artificial structure can represent the natural monomeric Ups1/Mdm35 structure.

**Figure 1.**
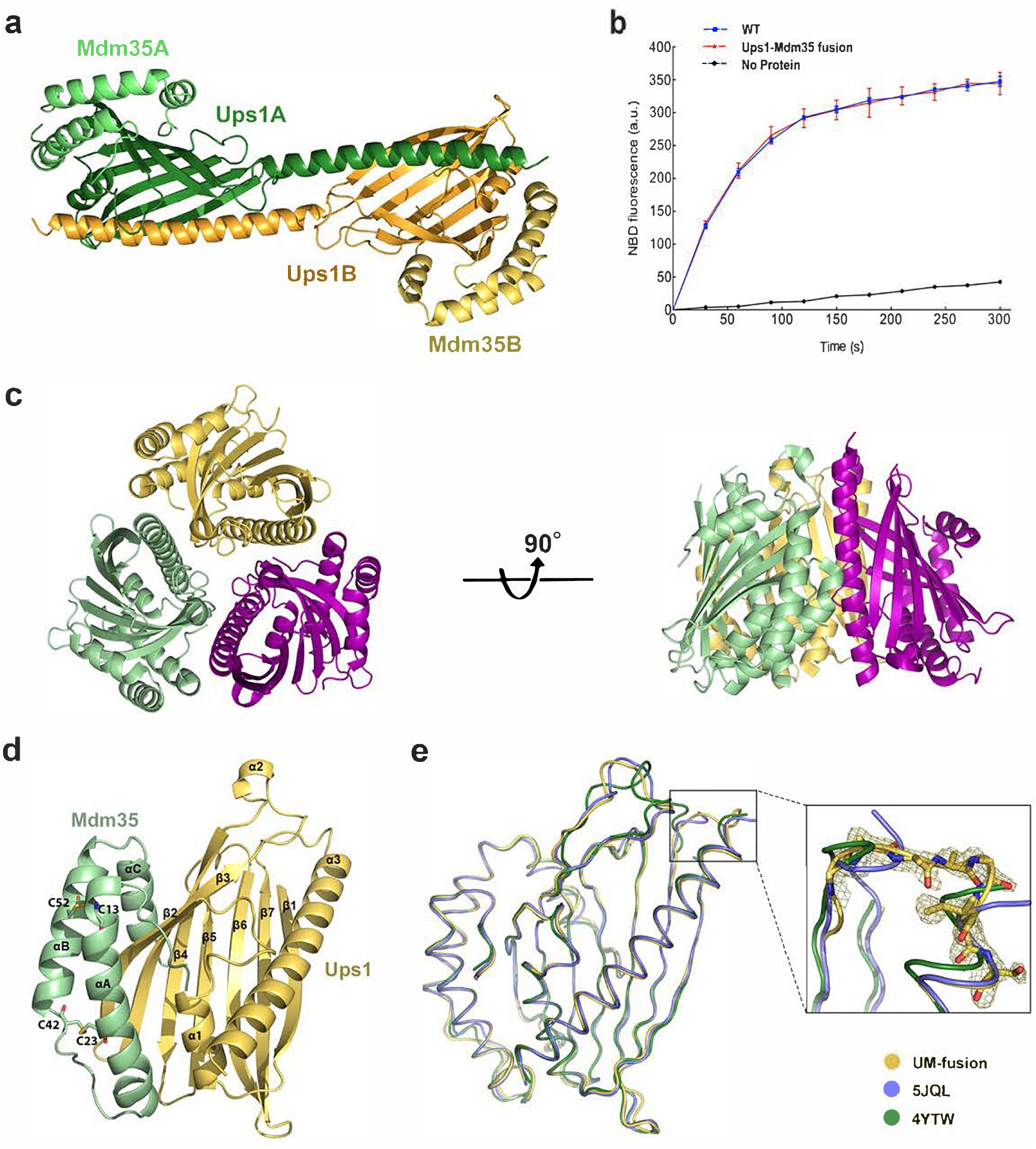
Crystal structures of Ups1/Mdm35 complex and Ups1-Mdm35-fusion protein. (**a**) The domain-swapped dimer of Ups1/Mdm35 (PDB code 5JQL). Ups1A, Ups1B, Mdm35A and Mdm35B are colored in forest, orange, light green and light yellow, respectively. (**b**) PA transfer activities of Ups1/Mdm35 and Ups1-Mdm35 fusion protein. Traces show means ± s.d. of three independent experiments. (**c**) Structure of the trimeric Ups1-Mdm35 fusion proteins (UM-fusion) in an asymmetric unit (PDB code 5JQM). (**d**) Ribbon diagram of UM-fusion. The Ups1 part is shown in light yellow and the Mdm35 part is shown in light green. The secondary structures are labelled accordingly. (**e**) Superposition of Ups1/Mdm35 structures (PDB codes 5JQL and 4YTW) and UM-fusion structure. Magnified view shows the electron density map of the L8 Loop of Ups1 in UM-fusion structure.

**Figure 2.**
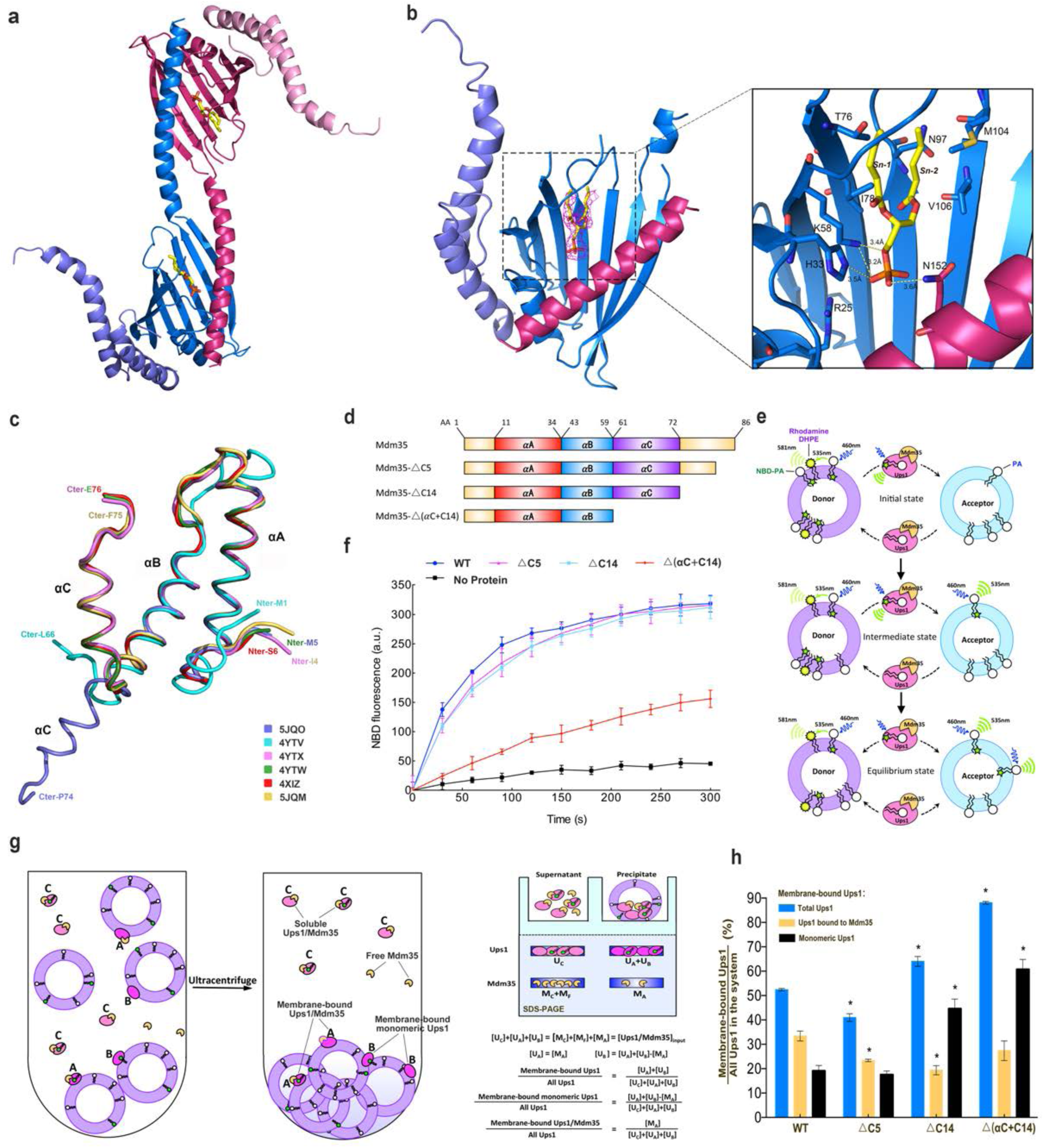
Structure of Ups1/Mdm35-DHPA, PA transfer activity and lipid co-sedimentation assay. (**a**) The domain-swapped dimer of Ups1/Mdm35-DHPA in an asymmetric unit (PDB code 5JQO). Ups1 is colored in hot pink and blue. Mdm35 is colored in slate and pink. The DHPA molecule is shown in stick with carbon atoms colored in yellow, oxygen atoms in red and phosphor atoms in orange. (**b**) Binding of DHPA in the pocket. The simulated annealing Fo–Fc difference map was calculated by omitting the DHPA molecule, and is shown with hot pink meshes countered at 3.0 σ. Magnified view shows the detailed interaction around the DHPA molecule. Broken lines designate possible hydrogen bonds. (**c**) Superposition of Mdm35 structures with the corresponding PDB codes annotated. (**d**) Schematic representation of Mdm35 and its truncation mutants. (**e**) Schematic diagram of the fluorescent-based PA transfer assay. (**f**) PA transfer activities of the WT and mutants. (**g**) Schematic diagram of liposome co-sedimentation assay. The formula is used to quantify the percent of different state of Ups1 binding to liposome. (**h**) Quantification of liposome co-sedimentation assays. Data are representative of three independent experiments. All results are expressed as mean values ± standard deviation (SD). Independent samples t-test was used for statistical analyses by SPSS 23.0 (n=3), (‘*’*p*<0.01).

To get a monomeric structure of Ups1/Mdm35, we designed an Ups1-Mdm35 fusion protein, in which a PreScission protease recognition sequence LEVLFQGP was inserted between the C-terminus of Ups1 and the N-terminus of Mdm35. This fusion protein has the same PA transfer activity as the wild type (Fig.1b). A similar phenomenon was observed for another single chain Ups1-Mdm35 fusion protein by Y. Watanabe et al ^17^. The structure of Ups1-Mdm35 fusion protein was solved (PDB code 5JQM) and refined to 1.5 Å by molecular replacement. There are three monomeric Ups1-Mdm35 fusion proteins per asymmetric unit (Fig.1c). The linker between Ups1 and Mdm35 is invisible suggesting its high flexibility. Because the three molecules superimpose well with the RMSD (root mean square of deviation) of ∼ 0.1 Å, we chose the most complete Ups1 model from chain B and Mdm35 model from chain C to rebuild the structure of Ups1-Mdm35 fusion protein (referred as UM-fusion in the following context) for further structural analysis (Fig.1d).

In the UM-fusion structure, the monomeric Ups1 forms a “hot-dog” fold by seven-stranded antiparallel β-sheets (β1-β7) and three α-helices (α1-α3). Residues between β3 and β4 form the α2-loop including the α2-helix (Fig.1d). The overall structure of UM-fusion is similar to those of the artificial Ups1/Mdm35 monomers (PDB codes 5JQL and 4YTW), with RMSD of 0.704 and 0.766 Å for 239 and 230 Cα atoms, respectively (Fig.1e). The biggest difference is that electron densities of residues F133-K138 in UM-fusion are clear and continuous, but are rather weak in artificial monomers due to domain swap. Mdm35 contains three α-helices (αA-αC) with two disulfide bridges (C13-C52 and C23-C42) between αA and αB-helices (Fig.1d). These three α-helices embrace Ups1 to give a buried surface area of 1,222 Å^2^, which is about 12.7% of the whole surface area of Ups1. The PA binding pocket of Ups1 and the interactions between Ups1 and Mdm35 have been described in details by previous reports ^17, 18^.

### Structure of Ups1/Mdm35 bound with DHPA

To investigate the PA transfer mechanism of Ups1/Mdm35, we attempted to get the structure of PA-bound Ups1/Mdm35. Both Ups1/Mdm35 and Ups1-Mdm35-fused protein were co-crystallized with PA with different acyl chains, including DOPA (1,2-dioleoyl-sn-glycero-3-phosphate, 18:1–18:1), POPA (1-palmitoyl-2-oleoyl-sn-glycero-3-phosphate, 16:0–18:1) and DHPA (1,2-dihexanoyl-sn-glycero-3-phosphate, 6:0–6:0). Although we could obtain crystals of Ups1/Mdm35 with DOPA and DHPA, only crystals of the Ups1/Mdm35 with DHPA could diffract well and be suitable for structure determination. Finally, we determined the structure of Ups1/Mdm35-DHPA complex (PBD code 5JQO) at 3.6 Å resolution by molecular replacement with the UM-fusion protein (PBD code 5JQM) as the starting model. There are two Ups1/Mdm35 molecules per asymmetric unit (Fig.2a). The Mdm35 molecule presents an unprecedented conformation, in which its αC-helix does not bend towards the N-terminal of the αB-helix, but extends in the direction of the αB-helix and does not interact with Ups1 (Figs. 2a and 2c).

The two Ups1 molecules form a domain-swapped dimer as in Ups1/Mdm35-DLPA structure (PDB code 4YTX). To be noted that, F. Yu et al reported a monomeric Ups1/Mdm35-PA structure (PDB code 4XHR) ^18^. However, when we revisited this structure and found residues N134-I137 poorly fit with the density map (Supplementary Fig. 3), we rebuilt this part of structure, which turned out that Ups1 in this structure (PDB code 4XHR) is also domain-swapped.

In the crystal structure of Ups1/Mdm35-DHPA, each Ups1/Mdm35 contains a DHPA molecule with high RSCC (real space correlation coefficient) (Supplementary Fig. 4). Structural comparison of the Ups1/Mdm35-DHPA and Ups1/Mdm35-DLPA shows a similar Ups1 conformation with RMSD of 0.86 Å for 155 Cα atoms (Supplementary Fig. 5). In the structure of Ups1/Mdm35-DHPA, residues V66-G72 at the α2-loop could not be assigned due to lack of electron density, suggesting the high flexibility of this loop. The phosphate head of the bound DHPA is close to the positively charged side chain of R25, and the side chains of H33, K58 and N152 are within the distances that allow formation of electrostatic interactions or hydrogen bonds with the phosphate oxygen atoms. The sn-1 and sn-2 acyl chains of DHPA are stabilized via hydrophobic interaction with T76, I78, N97, M104 and V106 (Fig.2b). By comparison with the Ups1/Mdm35-DLPA structure, we found DLPA inserts into the positively charged pocket more deeply than DHPA (Supplementary Fig.6). This may be due to the acyl chain of DLPA is longer than that of DHPA, so that DLPA needs to occupy more space in the pocket.

### A new interaction mode between Ups1 and Mdm35

In the previously reported Ups1/Mdm35 structures, all three α helices of Mdm35 interact with Ups1. However, in the structure of Ups1/Mdm35-DHPA, only αA and αB helices of Mdm35 interact with Ups1, giving a buried surface area of about 790 Å^2^, which is about 8.1% of the whole surface area of Ups1. This is about 65% of the buried surface area between Ups1 and Mdm35 in other Ups1/Mdm35 structures. Thus, by changing the orientation of the αC helix, Mdm35 can adjust its interaction strength with Ups1.

To examine whether the interaction strength between Ups1 and Mdm35 would affect the PA transfer activity, we made three Ups1/Mdm35 truncation mutants (Fig.2d) for functional assays. The △C5 mutant lacks the C-terminal 5 residues (K82-K86) of Mdm35. The △C14 mutant lacks the C-terminal 14 residues (A73-K86), which are located just after the αC helix and untraced in the crystal structures. Secondary structure prediction shows that this segment is disordered. In the △(αC+C14) mutant, the αC helix of Mdm35 was further deleted, as a result this mutant can only bind to Ups1 in a weak interaction. First, we measured the PA transfer activities of these mutants by fluorescent-based PA transfer assay (Fig.2e). The △C5 mutant was used as a negative control, since Watanabe et al had proved this mutant has no effect on the PA transfer activity of Ups1/Mdm35 ^17^. As shown in Fig.2f, the △C5 and △C14 truncations have little effect on the NBD fluorescence increase, whereas the △(αC+C14) truncation dramatically impairs the PA transfer activity. This indicates the αC helix of Mdm35 plays an important role in PA transfer process and suggests that as the weakened interaction between Ups1 and Mdm35 decreases the PA transfer activity of Ups1/Mdm35.

We wanted to know whether the interaction strength between Ups1 and Mdm35 would affect the membrane binding ability of Ups1, thus the liposome co-sedimentation assay was performed (Fig.2g). When a system composed of Ups1/Mdm35 and PA-containing liposomes reaches chemical equilibrium, the following molecules should exist according to the published reports. First, there are liposome-bound Ups1/Mdm35 complex. This can be deduced from the facts that Mdm35 alone does not bind to liposomes, while liposome-bound Mdm35 was detected in Ups1/Mdm35 liposome flotation assay ^8, 21^. Second, there are membrane-bound monomeric Ups1, which were detected in liposome flotation assay ^8, 17, 21^. Third, there are cytosolic Ups1/Mdm35. Finally, there are cytosolic free Mdm35, which have dissociated from liposome-binding Ups1. The soluble proteins and the liposome-bound proteins can be separated by ultracentrifugation. The resulting supernatant and precipitate fractions were analyzed by SDS-PAGE. By quantifying the Ups1 and Mdm35 bands in the two fractions, the distribution of the two different states of Ups1 on liposomes can be calculated. We found that the overall membrane binding ability of Ups1 increases most significantly in △(αC+C14) mutant, compared to that of WT (Fig.2h). This is mainly due to the significant increase in the amount of membrane bound monomeric Ups1. This suggests that once the interaction between Ups1 and Mdm35 is weakened, more Ups1 monomer will tend to stay on the lipid membrane. Interestingly, we found the amount of membrane-bound monomeric Ups1 in △C14 mutant was also significantly increased compared to that of the WT. This suggests that the C-terminal 14 residues of Mdm35 are likely to be involved in the interaction with Ups1, although the high flexibility of this segment makes it invisible in the crystal structures.

### The interaction between Ups1/Mdm35 and membrane

By liposome co-sedimentation assay, we have detected that both Ups1/Mdm35 and monomeric Ups1 could bind to the membrane. To investigate how Ups1 interacts with the membranes in these two different states, we performed all atom molecular dynamics (MD) simulations in which either an Ups1/Mdm35 complex or a freestanding Ups1 (Ups1_free_) interacting with DOPA-containing lipid bilayers (Supplementary Movies 1 and 2). The initial orientation of Ups1 relative to the lipid membrane was defined according to the structural comparison between Ups1/Mdm35 and PITP-α. Because the α2-loop of Ups1 was thought to be the equivalent region of the lipid exchange loop of PITP-α, it may interact with the membrane in the same orientation as the lipid exchange loop. Thus we chose the initial orientation of Ups1 to make its α2-loop facing onto the membrane (Fig. 3a). The time evolution RMSD profiles show that, the conformational change of Ups1_free_ is relatively small and converges sufficiently after 300 ns, whereas those of Ups1/Mdm35 complex are larger for both Ups1 (Ups1_complex_) and Mdm35 parts and seem not to converge within the simulation time (Fig. 3b). Because Mdm35 was thought to stabilize residues at the interface between Ups1 and Mdm35, the increased RMSD of Ups1_complex_ may have hinted for allosteric sites at their interface. This speculation was confirmed by the root-mean-square-fluctuation (RMSF) profiles of Ups1 in these two simulations (Fig.3c). As expected, Ups1_complex_ has a significant lower residual fluctuation at the loop region (residues N43-N49) surrounded by Mdm35_complex_ (∼1.5 Å) than that in Ups1_free_ (∼2.5 Å), and had relatively higher RMSF at some distant segments (residues N28-H33, V66-L70, L100-G104 and K140-V163). The higher RMFS would affect the subsequent membrane binding of Ups1.

**Figure 3.**
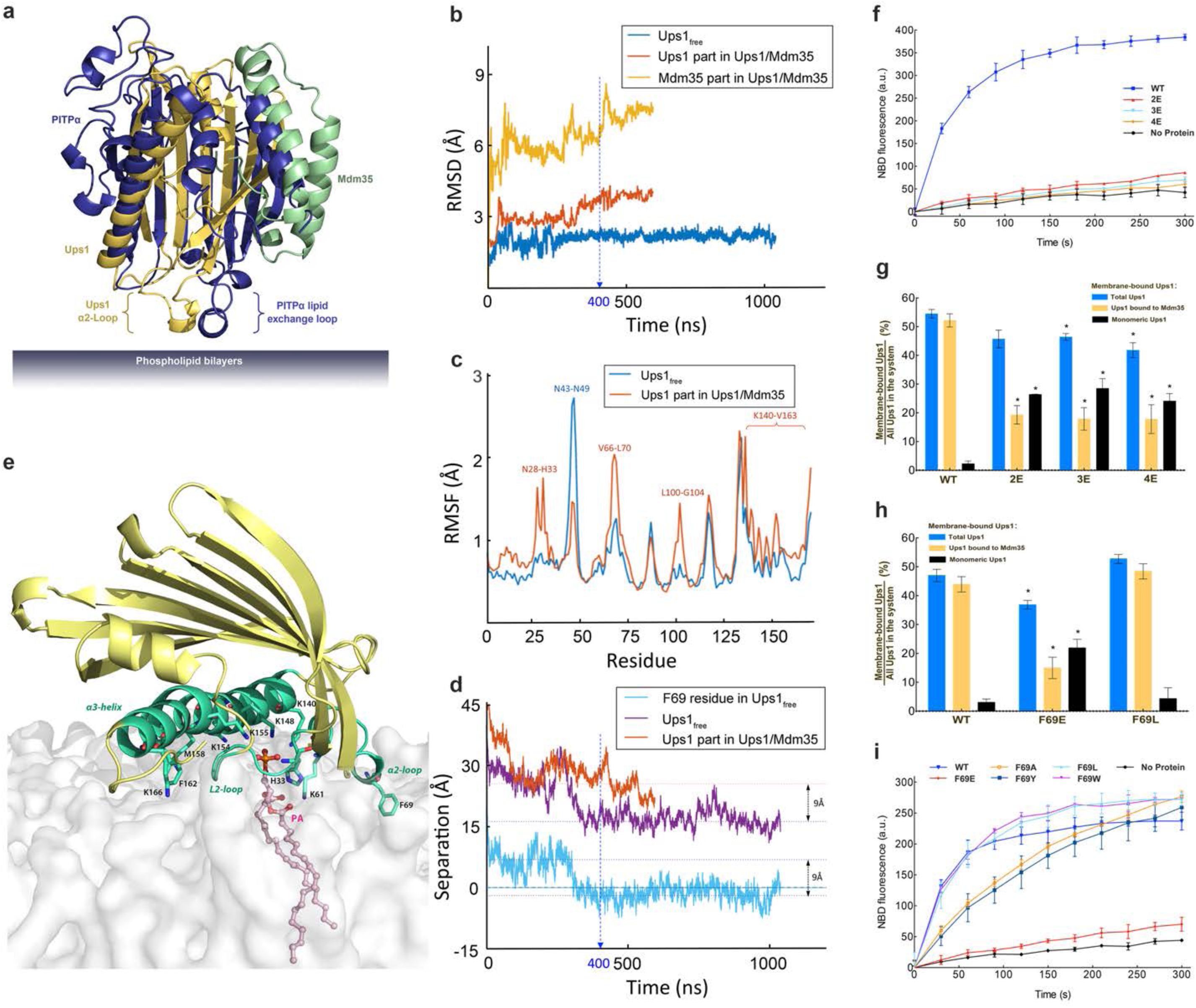
All-atom molecular dynamics simulations of Ups1/Mdm35 or Ups1_free_ interacting with the lipid bilayer. (**a**) Initial orientation of Ups1/Mdm35 or Ups1_free_ relative to lipid bilayer in MD simulations. The structure of Ups1/Mdm35 is superposed onto apo mouse PITPα (PDB code 1KCM). The identified lipid exchange loop in PITPα and the equivalent α2-loop in Ups1 are indicated. (**b**) RMSD profile of Ups1/Mdm35 and Ups1_free_ simulations. (**c**) RMSF profile of Ups1. RMSF was calculated for each Cα atom of Ups1 over the last 200 ns for both simulations. The trajectories were aligned with their first frame, as the reference structure, before calculating the average positions of the Cα atoms. (**d**) The time evolution profiles of the distances between the lipid phosphate plane and COM of Ups1 or F69 in Ups1_free_. After ∼ 400 ns, the relative distance between the COM of Ups1_free_ (or F69) and the membrane surface is decreased ∼9 Å. (**e**) Snapshot of Ups1_free_ interacting with the lipid bilayer at 1023 ns. Ups1_free_ is colored in yellow and the secondary structures involved in membrane binding (α2-loop, L2-Loop and α3-helix) are colored in lime green. The hydrophobic and positively charged residues involved in membrane binding are indicated. The PA molecule is shown in ball and stick with carbon atoms colored in yellow, oxygen atoms in red and phosphor atoms in orange. (**f**), (**i**) PA transfer activities of the wild type and mutants of Ups1/Mdm35. (**g**), (**h**) Quantification of liposome co-sedimentation assays. Data are representative of three independent experiments. All results are expressed as mean values ± standard deviation (SD). Independent samples t-test was used for statistical analyses by SPSS 23.0 (n=3), (‘*’*p*<0.01).

The time evolution profiles of the distances between the membrane surface phosphate plane (MSPP) and the center-of-mass (COM) of Ups1 in these two simulations show a sharp decrease of the distance for Ups1_free_. After 400 ns Ups1_free_ has a stable distance to the membrane, which is significantly different to Ups1_complex_ (Fig. 3d). This is also reflected by the converged RMSD profile of Ups1_free_ after 400 ns. And Ups1_free_ is found to establish stable protein-membrane interaction by inserting F69 that is located at the α2-loop into the lipid bilayers (Fig. 3d). The insertion of F69 into the membrane causes the COM of Ups1_free_ to approach the membrane bilayer by 9 Å, which should facilitate PA entering the lipid-binding pocket of Ups1. As expected, the direct interaction between PA and Ups1_free_ was observed after 800 ns (Supplementary Movie 3). A snapshot of Ups1_free_ interacting with the membrane at 1,023 ns is studied here (Fig. 3e), in which a PA molecule is trying to enter the lipid-binding pocket of Ups1_free_. The phosphate head of PA is surrounded by four positively charged residues H33, K61, K148 and K155, which had been determined important for the PA transfer ^17, 18^. Ups1_free_ interacts with the lipid membrane through the hydrophobic and positively charged residues located in the L2-loop (N28-H33), α2-loop (K61-R71) and the C-terminal long α3-helix (K140, R146, K154, M158, F162, K166 and R171). To be noted that, these membrane-bound structural elements contain exactly those residues with the high RMSF in Ups1_complex_ (Fig. 3c). The L2-loop is conserved, indicating its importance for Ups1 function (Supplementary Fig.7). The α2-loop is believed the lid for the PA binding pocket and may participate in membrane binding ^21^. As for the C-terminal long α3-helix, despite no significant sequence homology, its secondary structure is conserved among the members of PRELID family. Interestingly, we found although Ups1/Mdm35 does not establish a stable interaction with the membrane, it tries to approach the membrane in the same orientation as Ups1_free_ after 200 ns (Supplementary movie 1).

To test whether Ups1 interacts with the membrane as the MD simulation predicted, we performed the following experiments. First, we tested whether the C-terminal α3-helix of Ups1 is involved in membrane binding. Previous studies revealed that Ups1 exclusively binds to liposomes containing negatively charged phospholipids ^8^. And our MD simulation shows that a number of positively charged residues (K140, R146, K154, K166 and R171) in the α3-helix contact the negatively charged membrane surface. If these basic residues really participate in membrane binding, mutating them into Glu should impair the PA transfer activity of Ups1/Mdm35. We constructed a series of mutations on these residues and tested their abilities of membrane binding and PA transfer. The Ups1-2E mutant contains K140E and R146E double mutations. The Ups1-3E mutant contains K140E, R146E and K154E triple mutations. And the Ups1-4E mutant contains K140E, R146E, K154E and K166E quadruple mutations. We found Ups1-2E alone is enough to completely eliminate the PA transfer activity of Ups1 (Fig.3f). Since structural studies of Ups/Mdm35 had excluded the possibility of interaction of these basic residues with Mdm35 or PA, the loss of PA transfer activity in these mutants would be due to disruption of the Ups1-membrane interaction. This hypothesis was ascertained by the result of liposome co-sedimentation assay. As shown in Fig. 3g, for these mutants, the amount of membrane-bound Ups1/Mdm35 (defined as State A) decrease significantly, which leads to the decrease of their overall membrane binding abilities, compared to that of WT. It is worth noting that for these mutants the amount of membrane bound monomeric Ups1 (defined as State B) significantly increase, compared to that of WT, which will be discussed later.

Then we examined whether F69 is involved in membrane binding of Ups1/Mdm35. F69 is located in the α2-loop, which contains positively charged and hydrophobic residues, but no negatively charged residues. Its amino acid composition and structural similarity with the lipid exchange loop of PITP-α make it possible to participate in binding to the negatively charged membrane (Supplementary Fig.7). If F69 is involved in membrane binding, mutating F69 to the negatively charged residue Glu should impair the membrane binding ability of Ups1/Mdm35, whereas mutating F69 to a residue with a similar side chain, such as Leu, should not have much effect on its membrane binding ability. The result of liposome sedimentation assay confirmed our hypothesis (Fig.3h). The overall membrane binding ability of F69E mutant is significantly impaired. Meanwhile, F69E mutation also causes a large accumulation of monomeric Ups1 on the membrane. As for F69L mutant, the amount of Ups1/Mdm35 and monomeric Ups1 on the membrane is similar to that of WT. To test whether it is possible for F69 to be buried into the membrane during the PA transfer process, we further mutated F69 into small hydrophobic (Ala), bulky uncharged polar (Tyr) and bulky hydrophobic (Trp) side residues and measured their PA transfer activities (Fig.3i). The F69E mutant nearly abolished the PA transfer activity. Both F69A and F69Y mutants markedly impair the PA transfer activities. As for F69W and F69L mutants, they keep the same PA transfer activities with WT. Since F69 can only be replaced by residues with large hydrophobic residues, it should be in a completely hydrophobic environment at some stage at least during the PA transfer process. Structural analysis shows that F69 is unlikely to interact with the hydrophobic residues of Ups1 itself, thus a reasonable explanation is that this residue would be buried in the membrane during the PA transfer process, as it is predicted from MD simulation.

### Potential conformational change upon PA binding

To explore how PA enters the PA-binding pocket of Ups1 and whether Ups1 has a conformational change before and after PA binding, we compared the crystal structures of Ups1 in different states, including the MD simulated membrane-bound state (Ups1_free_-1023ns), the apo state (PDB codes 5JQM and 4YTW), the DLPA-bound (PDB code 4YTX, chain B) and DHPA-bound state (PDB code 5JQO). In addition, Y. Watanabe et al observed another state of Ups1 (PDB code 4YTX, chain L), in which Ups1 has a more open α2-loop, yet has no PA in its lipid binding pocket ^17^, which is referred to X-state here.

After superimposing the five crystal structures to the MD simulated membrane bound structure of Ups1, we found although the overall structures of the six molecules are very similar, the conformations of the α2-loop and the N-terminus of the α3-helix (N-α3-helix) are obviously different (Fig.4a). In the Ups1_free_-1023ns and apo state structures (PDB codes 5JQM and 4YTW), the hydrophobic side chains of P63 and W65 in the α2-loop bind to the hydrophobic region formed by I137, V141 and W144 of the N-α3-helix, giving a buried surface area of 499 Å^2^, 344 Å^2^ and 234 Å^2^ (∼31%, 24% and 15% of the total surface area of the α2-loop), respectively (Fig. 4b and Table 1). However, in the PA-bound and the X-state structures, both the α2-loop and N-α3-helix swing outward, which increases the distance between the α2-loop and N-α3-helix and destroys their hydrophobic interaction (Fig. 4a). To better compare the changes of proximity between the α2-loop and N-α3-helix in these structures, we selected four adjacent residues (P63, W65, V141 and W144) located in the α2-loop and N-α3-helix (Fig. 4b) and measured the distance between them (P63-V141, P63-W144, W65-V141, and W65-W144). As shown in Table 1, the four distances are the smallest in Ups1_free_-1023ns structure, followed by the apo-state, PA-bound state, and finally the X-state. Because both α2-loop and N-α3-helix are involved in membrane binding, the change of their proximity leads to the change of the opening size on the membrane interaction interface of Ups1 (Fig. 4c). For the membrane-anchored Ups1_free_-1023ns and the apo state structure, there is only a small hole on this interface, which is lined by residues H33, K61, K148 and K155 and might be involved in recognizing or binding the head of PA. For the two PA-bound Ups1 structures, the hole is enlarged into a cleft. We speculated it might be due to the PA occupation in the pocket. In the X-state, the cleft is continued to expand into a wedge-shaped cavity. Although there is no density of DLPA in its PA binding pocket, we speculated the structure might represent an intermediate state in which PA is entering or exiting from the pocket, thus requiring a wider channel on the membrane interaction interface to accommodate the two long acyl chains of PA. To test our hypothesis, we re-analyzed the electron density of the X-state and found an additional density near the membrane contact surface, which could accommodate a DLPA molecule well (Fig.4d).

**Figure 4.**
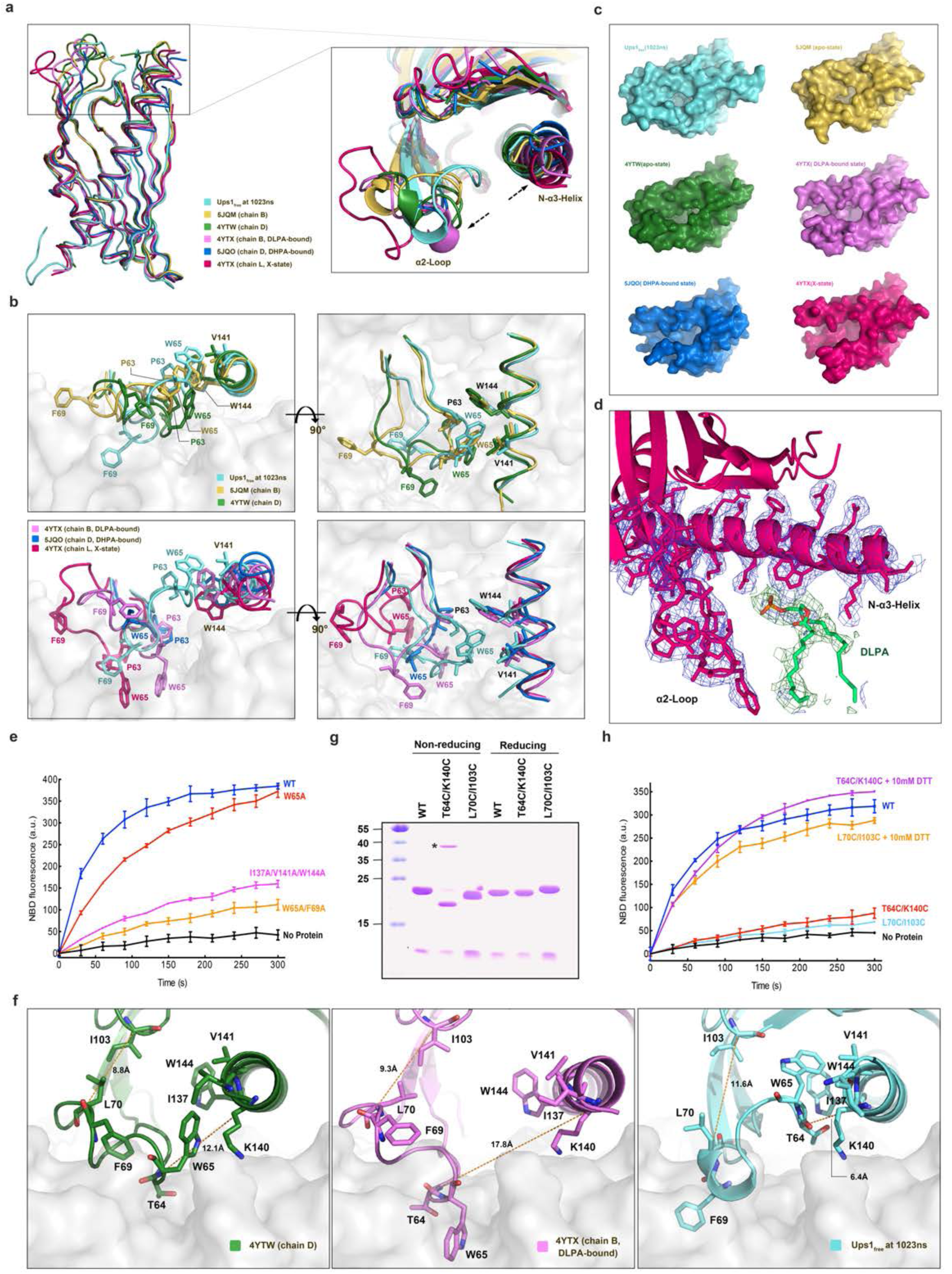
Conformational changes of Ups1/Mdm35 during PA entry and binding. (**a**) Superposition of six Ups1 structures with the corresponding PDB codes annotated. The magnified view shows the conformational changes of the α2-loop and N-α3-helix. (**b**) Magnified view of the regions around P63, W65, V141 and W144 in (a). (**c**) The membrane-contacting surface of different Ups1 structures. (**d**) Electron density map of a newly discovered DLPA molecule binding to the X-state Ups1 (PDB code 4YTX). The X-state Ups1 is colored in hot pink. The DLPA molecule is shown in stick with carbon atoms colored in yellow, oxygen atoms in red and phosphor atoms in orange. The electron density map of the α2-loop and N-α3-helix is colored in marine. The simulated annealing Fo–Fc difference map was calculated by omitting the DLPA molecule, and is shown with green meshes countered at 3.0 σ. (**e**) PA transfer activities of the wild type and mutants of Ups1/Mdm35. (**f**) Distances between the Cα atoms of L70 and I103 and between the Cα atoms of T64 and K140 in the corresponding Ups1 structures. (**g**) The T64C/K140C and L70C/I103C mutants were analyzed by SDS-PAGE with or without 10 mM DTT. (**h**) PA transfer activities of the wild type and mutants of Ups1/Mdm35. For the reducing condition, T64C/K140C and L70C/I103C mutants were pretreated with 10mM DTT before analysis.

**Table 1.** Comparison of the distances between the α2-loop and N-α3-helix of Ups1 in different states.

| State | Source | P63-V141 | P63-W144 | W65-V141 | W65-W144 | Interaction area* |
| --- | --- | --- | --- | --- | --- | --- |
| Membrane-anchored state | Simulated structure of Ups1<sub>free</sub> at 1023ns | 8.5 | 7.2 | 6.5 | 8.3 | 499 |
| Apo-State | 5JQM (chain B) | 8.6 | 7.9 | 8.8 | 10.5 | 350 |
|  | 4YTW (chain D) | 10.2 | 8.2 | 11.3 | 11.2 | 234 |
| PA-bound State | 4YTX (chain B, DLPA) | 14.4 | 12.9 | 15.3 | 15.1 | 0 |
|  | 5JQO (chain D, DHPA) | 15.1 | 14.3 | 16.8 | 18.1 | 0 |
| X-State | 4YTX (chain L, ?) | 20.3 | 19.2 | 20.9 | 21.0 | 0 |
| The unit of distance is Å. The unit of interaction area is Å<sup>2</sup>. * The interaction area is between the α2-loop (K61-I73) and the N-α3-helix (G136-T147) of Ups1. | | | | | | |

The structural comparison further reveals that the positions of F69 and W65 vary among different states (Fig.4b). In PA-bound and X-state structures, it is W65 that is inserted into the membrane to anchor Ups1. In Ups1_free_-1023ns structure, F69 is inserted into the membrane. In apo-state structures, both F69 and W65 are not inserted into the membrane. It seems that the conformational changes of the α2-loop and N-α3-helix result in the membrane-anchoring residue of Ups1 alternating between W65 and F69 during the PA transfer process (Supplementary movie 4).

To test whether the hydrophobic interaction between the α2-loop and N-α3-helix will affect the function of Ups1/Mdm35, we generated a W65A mutant and an I137A/V141A/W144A triple mutant of Ups1. Both of them have an impaired PA transfer activity (Fig.4e). We also constructed a W65A/F69A double mutant and found the PA transfer activity of Ups1 is nearly abolished (Fig.4e).

To test whether the alternate insertion of F69 and W65 into the membrane is essential for the function of Ups1/Mdm35, we wanted to lock the position of W65 or F69 so that the membrane anchoring of Ups1 can only be accomplished by one of them (Fig.4f). First, we designed the L70C/I103C mutant in which F69 cannot be inserted into the membrane. In the PA-bound structures, W65 interacts with I137, V141 and W144, and F69 is located above the membrane surface. The distance between the Cα atoms of L70 and I103 in the DLPA-bound structure (PDB code 4YTX, chain B) is 8.8 Å. If a disulfide bond is formed between L70 and I103 by mutating to cysteines, the distance between their Cα atoms should be reduced to 5.5-5.7 Å, which should make the reside (L70C) away from the membrane. As a result, the adjacent residue F69 should also be away from the membrane. Meanwhile, it should be less affected for W65 that still has a chance to insert into the membrane. Second, we designed another T64C/K140C mutant based on the structure of Ups1free-1023ns, in which W65 cannot be inserted into the membrane (Fig.4f). The position of W65 is locked to interact with I137, V141 and W144 in this mutant while F69 is almost unaffected and still has the ability to insert into the membrane.

After purifying the two mutants, we found that each pair of the introduced cysteines can form an intramolecular disulfide bond spontaneously as expected. As shown in Fig. 4g, the bands of L70C/I103C and T64C/K140C mutants shift upward upon DTT treatment. Furthermore, the disulfide bonds between L70C and I103C, and between T64C and K140C were validated by mass spectrum (Supplementary Fig. 8). Although these two intra-cross-linked mutants could hardly transfer PA, their PA transfer activities were restored after 10 mM DTT treatment (Fig.4h). These data suggest that the conformational changes (membrane insertion) of both F69 and W65 are important for PA transfer activity of Ups1/Mdm35. Interestingly, for the T64C/K140C mutant, we also detected a small amount of inter-cross-linked domain-swapped dimer, which was not detected in the previous N28C /G159C mutant ^17^.

### Regulation of PA transfer activity by pH

Because the pH of IMS is significantly influenced by the respiratory chain function ^25^, we guessed that the pH value would regulate the PA transfer activity of Ups1/Mdm35. After measuring the PA transfer activity of Ups1/Mdm35 under different pH, we found that the optimum pH of Ups1/Mdm35 is 7.0 (Fig. 5a). At pH 6.5 and pH 7.5, the activity is slightly lower. At pH 6.0, the activity is significantly lower than that at pH 7.0. And at pH 5.5, the activity is even less and falls to less than one-third. Considering there are many basic residues located in the membrane binding surface of Ups1, we suspect that the pH may affect the affinity between Ups1/Mdm35 and the membrane by changing the charge on these residues. To test this, the electrostatic potentials and net charge of Ups1 molecules in different states under different pH values were calculated (Table 2) using the PDB2PQR server ^26^. We found all of the molecules carry the largest number of positive charges at pH 5.5 and the number of charges carried by Ups1 varies according to different states and pH. The change in electrostatic potentials under different pH mainly occurs at the entrance of the PA binding pocket (Fig. 5b). It seems that under lower pH, Ups1 would have a stronger membrane binding ability due to its increased positively charge. This hypothesis was confirmed by the result of liposome co-sedimentation assay (Fig.5c) that at lower pH more monomeric Ups1 accumulates on the membrane, which would be the reason for its decreased PA transfer activity (see discussions below).

**Figure 5.**
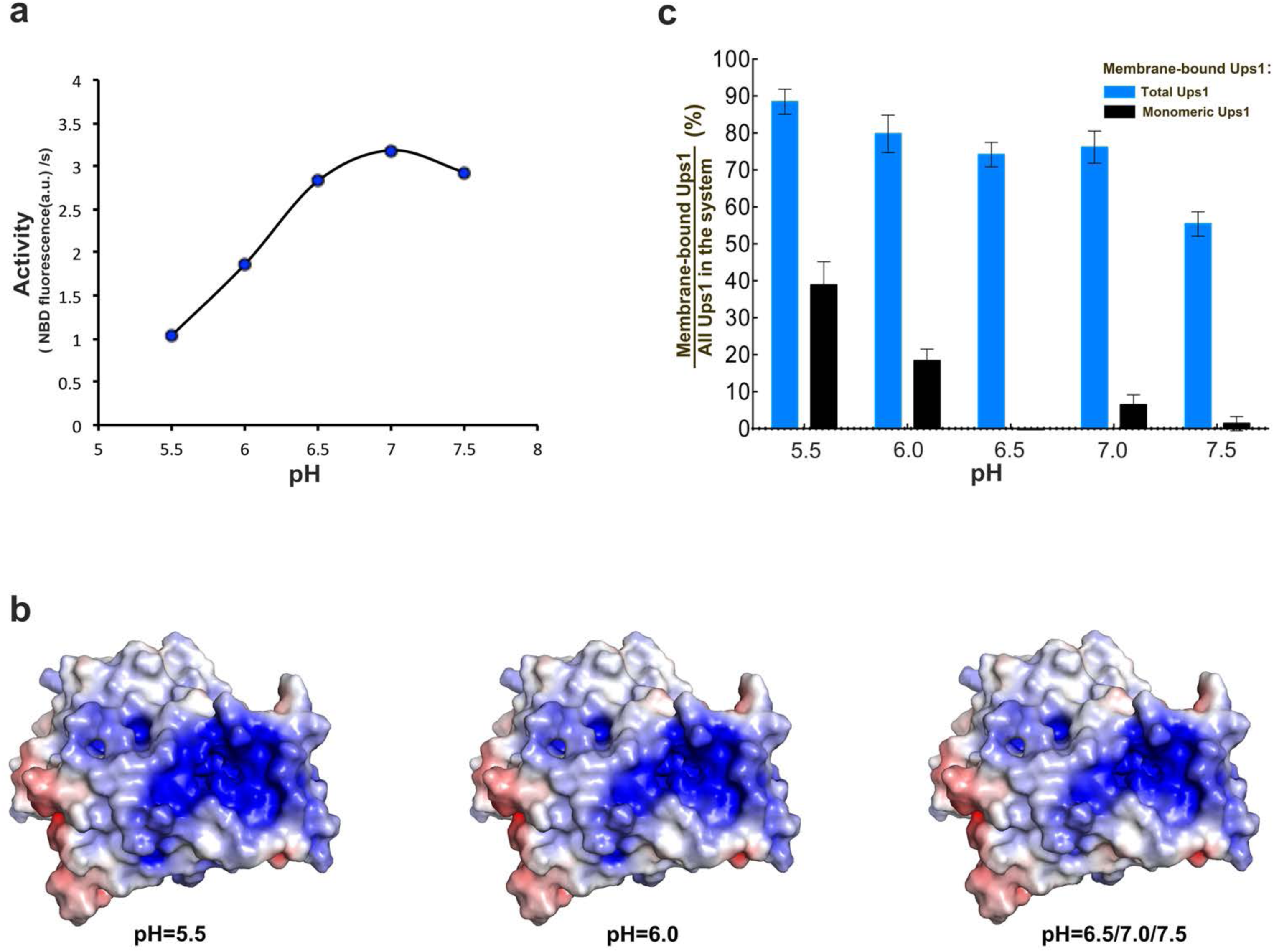
Effect of pH values on the PA transfer activity of Ups1/Mdm35. (**a**) pH dependent PA transfer activity of Ups1/Mdm35. For each reach, the fluorescence data from the first 60s was used to calculate the initial transfer rate of PA. (**b**) Surface electrostatic potentials of Ups1/Mdm35 under different pH. The change mainly occurs at the entrance of the PA binding pocket. (**c**) Quantification of liposome co-sedimentation assays at different pH. Data are representative of three independent experiments. All results are expressed as mean values ± standard deviation (SD). Independent samples t-test was used for statistical analyses by SPSS 23.0 (n=3), (‘*’*p*<0.01).

**Table 2.** The number of net charges carried by Ups1 under different pH.

|  | pH5.5 | pH6.0 | pH6.5 | pH7.0 | pH7.5 |
| --- | --- | --- | --- | --- | --- |
| 5JQM (Apo state) | 12 | 8 | 8 | 8 | 8 |
| 4YTW (Apo state) | 11 | 9 | 9 | 8 | 7 |
| Ups1<sub>free</sub>_1023ns (membrane-anchored state) | 11 | 10 | 8 | 8 | 8 |
| 4YTX (X-state) | 10 | 9 | 7 | 7 | 7 |
| 4YTX (DLPA-bound state) | 9 | 8 | 7 | 6 | 6 |
| Only Ups1 parts of these structure were used for calculation. The net charge was calculated by PDB2PQR Server.. |  |  |  |  |  |

## DISCUSSIONS

Although there have been many crystal structures of Ups1/Mdm35 (or its homologous) in different states reported ^17, 18, 21, 23^, the molecular details of how PA is transferred between membranes is yet unknown. In the present study, we reported another two crystal structures of Ups1/Mdm35 in a truly monomeric apo state and a novel DHPA bound state, which provides us an opportunity to compare all available structures systematically and gain further insights into the conformational changes of Ups1/Mdm35.

Besides, we developed the liposome co-sedimentation method to quantitatively detect membrane bound Ups1 in Mdm35 complexed and none-complexed free states, which discovered the relationship between the portion of membrane bound Ups1 in free state and the PA transfer activity of Ups1/Mdm35. Our dedicative all-atom molecular dynamics simulations further indicated that the none-complexed Ups1 can form a more stable interaction with membrane in comparison with Ups1/Mdm35. MD simulations further disclosed novel key elements, the α2-loop, α3-helix and L2-loop, involved in membrane binding and potential PA entry, which were subsequently verified by mutagenesis experiments. By comparing the PA transfer activities among different truncations of Mdm35, we found the interaction strength between Mdm35 and Ups1 also regulates the PA transfer activity of Ups1/Mdm35.

Based on all available crystal structures, MD simulations and all biophysical assays in this study, we could attribute different structures of Ups1/Mdm35 into membrane-free apo state (PDB codes, 5JQM and 4YTW), membrane-bound apo state (MD simulated structure), membrane-bound PA entry state (PDB code, 4YTX) and PA bound state (PDB codes, 5JQO and 4YTX). Both liposome co-sedimentation experiments and MD simulations indicated the none-complexed Ups1 is the important transit during Ups1/Mdm35 mediated PA transfer, which was also suggested previously that the extraction of PA from membrane or delivery of PA into membrane is performed by Ups1 independently ^8^. With these basic concepts as well as the key elements for PA transfer discovered in this study, we could propose a detailed molecular model for Ups1/Mdm35 mediated PA transport (Fig.6 and Supplementary Movie 4).

**Figure 6.**
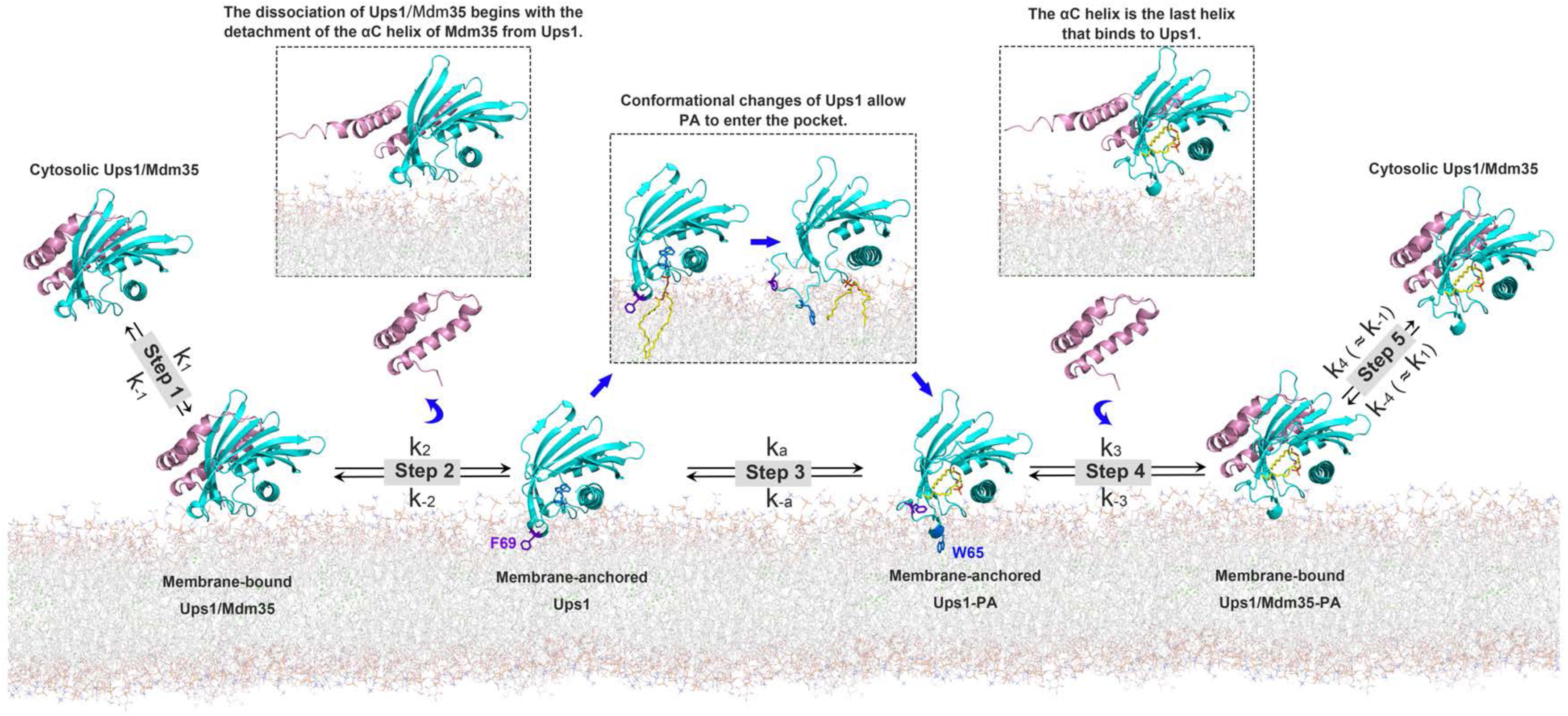
A working model for Ups1/Mdm35 mediated PA transport. Step 1, Ups1/Mdm35 interacts with the membrane via the membrane-binding residues of Ups1. Step 2, the dissociation of Mdm35 from Ups1 leads to the insertion of F69 into the membrane, anchoring Ups1 onto membrane. Step 3, the PA entrance and binding leads to the conformational change of Ups1, in which W65 is inserted into the membrane as a substitute of F69. Step 4, Mdm35 binds to Ups1-PA. Step 5, Ups1-PA /Mdm35 releases from membrane to solution.

First, when Ups1/Mdm35 approaches the lipid bilayers, Ups1/Mdm35 intends to interact with membrane through the membrane-binding residues of Ups1. These residues include N28-H33 of the L2-loop, and the hydrophobic and positively charged residues located in the α2-loop and the C-terminal long α3-helix. Although these membrane-binding residues are highly flexible in the presence of Mdm35, they can still interact with the membrane and allow Ups1/Mdm35 associated onto the membrane as an integral part (membrane-bound Ups1/Mdm35).

Second, Mdm35 will dissociate from Ups1, which begins with the αC-helix detachment, followed by the detachment of αA and αB helices. This liberates the residues of Ups1 located at the Ups1-Mdm35 interface and the resulting allosteric effects alleviate the flexibility of the membrane-binding residues of Ups1. Under this circumstance, the hydrophobic interaction (mediated by P63, W65, I137, V141 and W144) between the α2-loop and the N-α3-helix is enhanced, resulting in the insertion of F69 into the membrane. This event makes the COM of Ups1 further approaching to the lipid bilayer (membrane-anchored Ups1), which is important for PA extraction.

Third, the negatively charged phosphate head of PA is attracted to the positive charged residues H33, K61, K148 and K155, and interacts with them, which triggers conformational changes of the N-α3-helix and α2-loop. They move in opposite directions to enlarge the entrance of the PA-binding pocket, allowing the two long acyl tails of PA to enter into the pocket. During this process W65 is inserted into the membrane as a substitute of F69. When PA enters into the binding pocket completely, the N-α3-helix swings inwards a little to avoid the exit of PA from the pocket.

Fourth, Mdm35 interacts again with PA bound Ups1 to form Ups1/Mdm35-PA. In the fifth and final step, Ups1/Mdm35-PA releases from the membrane to the solution. The PA release process by Ups1/Mdm35 is just a reverse process of PA extraction and will not be discussed further.

This model can well explain the contradictory views in the literature concerning whether the α2-loop participates in membrane binding. Y. Watanabe et al reported the △lid mutant, lacking the α2-loop (lacking residues L62-R71), has an impaired PA transfer activity, whereas is normal in binding to membrane ^17^. They thought the α2-loop is not involved in membrane binding, but only plays as a gate for PA entry and exit. However, considering PA comes from the membrane, it’s difficult to imagine how α2-loop acts as a gate without membrane interaction.

In our model, the deletion of α2-loop does not abolish membrane binding of Ups1 but affects PA extraction ability, thus impairing PA transfer activity of Ups1/Mdm35.

Based on our model and the previously published kinetic model for phospholipid binding and transfer by the nonspecific lipid-transfer protein ^27^, the kinetic process of PA transfer by Ups1/Mdm35 can be described as the following reactions.

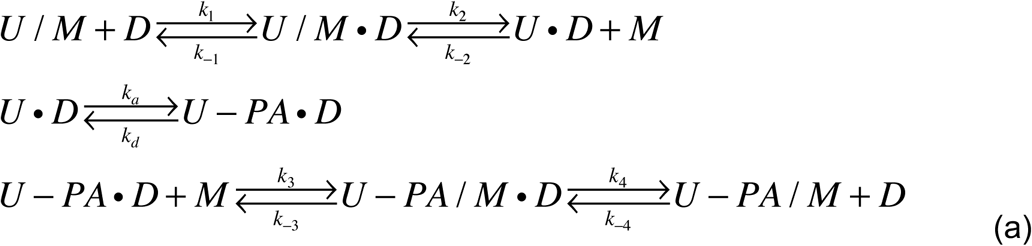

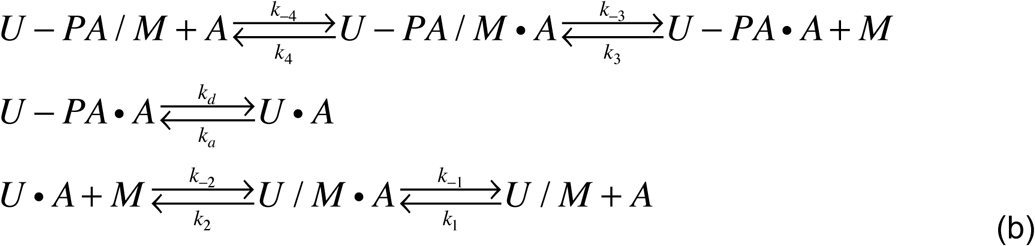

where U, M, U/M, D, A and PA represent Ups1, Mdm35, Ups1/Mdm35, donor membrane, acceptor membrane and phosphatidic acid, respectively. U-PA and U-PA/UM represent PA-bound Ups1 and PA-bound Ups1/Mdm35. U/M·D, U/M ·A, U-PA /M·D and U-PA/M·A represent membrane bound complexes. The forward/reverse reaction constants for each step are described as k_1_/k_-1_, k_2_/k_-2_, k_3_/k_-3_, and k_4_/k_-4_, and the association/dissociation constants of PA to Ups1 are denoted as k_a_/k_d_. The reaction (a) represents the process that Ups1/Mdm35 binds onto donor membrane and performs PA extraction while the reaction (b) represents the process that Ups1/Mdm35 binds to acceptor membrane and performs PA release. The reaction (b) is just a reverse procedure of the reaction (a), thus sharing the same reaction constants. In the following discussion, we assume that the association/dissociation reaction rates between U/M and membrane are similar with that between U-PA /M and membrane. Thus, k_4_/k_-4_ approximately equal to k_-1_/k_1_.

If steady state conditions are reached, ten equations can be derived from above reactions (see Supplementary Text). In liposome co-sedimentation experiments, the donor and acceptor membrane are identical. Thus, the equations in Supplementary Text can be simplified and the equilibrium constant (K1) of the binding between Ups1/Mdm35 and membrane can be derived as,

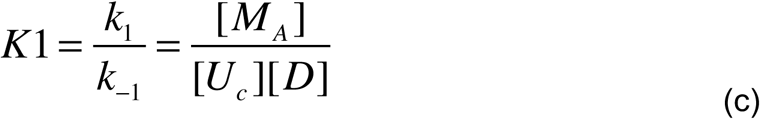

where [M_A_] represents the concentration of Mdm35 bound to Ups1 on the membrane, [U_C_] represents the concentration of Ups1 in the solution, and [D] represents the concentration of membrane (see Fig. 2g).

Then we could calculate the binding constant between Ups1/Mdm35 and membrane by utilizing the experimental data from liposome co-sedimentation (Supplementary Fig.9). We found the K1 values of 2E, 3E and 4E mutants are much lower than that of WT, indicating that mutating the positively charged residues at Ups1-membrane binding interface to negatively charged residues decreases the membrane binding ability of Ups1/Mdm35, which is in consistency with our prediction. Furthermore, the K1 value of △(αC+C14) mutant is much higher than that of WT, suggesting the mutant binds to the membrane more easily, which is also in consistency with our prediction that Ups1 needs to disassociate from Mdm35 and form a stable interaction with membrane, and the weaker binding between Mdm35 and Ups1 decreases the energy barrier of disassociation and increases the possibility for Ups1 binding to membrane. As a result, the above model of the detailed PA transfer by Ups1/Mdm35 is fairly consistent with our experimental data.

From the liposome co-sedimentation experiments and the PA transfer assays, we could found that weakening the interaction between Mdm35 and Ups1 increases the amount of total membrane bound Ups1, which is majorly contributed from the amount of monomeric membrane bound Ups1 (Fig. 2h). Besides, alteration of Ups1-membrane binding interface not only decreases the amount of membrane bound Mdm35/Ups1 complex but also increases the amount of monomeric membrane bound Ups1 (Figs. 3g and 3h). We could further found that the amount of monomeric membrane-bound Ups1 is inversely correlated with the overall PA transfer activity of Mdm35/Ups1 (Figs. 2f, 3f and 3i). According to our above molecular model of PA transfer, we could give a rational explanation of this inverse correlation. The success of PA transfer not only needs the binding of Ups1 to the membrane but also needs the successful release of Ups1 from the membrane. The weakened interaction between Ups1 and Mdm35 increases the energy barrier of releasing PA-bound Ups1 from the membrane, which is aided by Mdm35. Thus, we could observe more portion of monomeric Ups1 on the membrane. As a result, the overall PA transfer activity is decreased. The alteration of Ups1-membrane binding interface not only decreases the membrane binding ability of Ups1/Mdm35 but also could affect negatively the efficiency of PA entry and binding (to be noted that the PA entry site is coupled with the Ups1 membrane binding site), thus most monomeric Ups1 are trapped on the membrane without PA binding. As a result, the overall PA transfer activity is nearly eliminated. The inverse correlation between the amount of monomeric membrane bound Ups1 and the PA transfer activity could also be observed from the pH dependent experiments (Fig. 5).

The regulation of pH value to the PA transfer activity of Ups1/Mdm35 is very likely one of regulating mechanisms *in vivo*. In mitochondrion, protons are pumped into IMS, which would lower the pH value of IMS. Since CL is important for the assembly and stability of the respiratory chain complexes, lower pH suggests the respiratory chain functions well and there is enough CL to contribute. Therefore, the PA transfer activity of Ups1/Mdm35 needs to be reduced to decrease the synthesis of CL at IMM. In another case, an increased pH of IMS suggests a potential decreased activity of the respiratory chain, potentially due to the limited amount of CL. As a result, the higher PA transfer activity of Ups1/Mdm35 would be important to balance the synthesis of CL at IMM.

Overall, our present structural and biophysical studies give a deep insight to understand the PA transfer mechanism by Ups1/Mdm35, which provides additional knowledge of mitochondrial physiological homeostasis.

## MATERIALS AND METHODS

### Molecular cloning, protein expression and purification

To construct the co-expression vectors for N-terminal 6×His tagged full-length Ups1 and Mdm35, the corresponding genes were amplified by PCR from *S.cerevisiae* genome and cloned into the plasmid pETDute-1 (Novagen). To construct N-terminal 6×His tagged Ups1-Mdm35 fusion protein, the recognition sequence of PreScission protease (LEVLFQGP) was inserted between the C-terminus of Ups1 and the N-terminus of Mdm35 to produce an Ups1-Mdm35 fusion gene, which was also inserted into the plasmid pETDuet-1. The constructs containing point mutations were generated by PCR-based sitedirected mutagenesis. All constructs were sequenced to confirm their identities.

The target proteins were expressed in *E.coli* Trans B cells (TransGen Biotech) that were cultured in LB medium. After addition of 0.1 mM isopropyl-D-thiogalactoside (IPTG), the cells were cultured at 30 °C for 5 hr. Cells were collected and re-suspended in buffer A (50 mM Tris-HCl pH8.0, 300 mM NaCl, 20 mM imidazole, and 1 mM PMSF) and then disrupted using the high-pressure method at 120 MPa. Cell lysate was centrifuged with a JA25.5 rotor (Beckman Coulter) at 39,000 g for 40min at 4 °C. The supernatant was loaded onto a column of Ni-chelated Sepharose 6 Fast Flow (GE Healthcare) that was pre-equilibrated with buffer A. The target protein was eluted using buffer B (50 mM Tris-HCl pH8.0, 300 mM NaCl, and 150 mM imidazole) and concentrated by ultrafiltration (30 kDa cutoff; Amicon Ultra). The elution was further purified by size exclusion chromatography with Superdex75 (10/300) column (GE Healthcare) in buffer C (20 mM Tris-HCl, pH7.5 and 150 mM NaCl). The peak fractions were collected, concentrated to ∼20 mg/mL and stored in −80 °C. for further experiments. For Ups1-Mdm35 fusion protein, the elution was purified with Superdex75 in Buffer D (20 mM HEPES, pH7.0, and 150 mM NaCl). Then the peak fractions were desalted and further purified by ion-exchange chromatography using 1-ml Resource S column (GE Healthcare) in buffer E (20 mM mM HEPES, pH7.0) and buffer F (20 mM mM HEPES, pH7.0, and 1M NaCl). Two peaks were obtained using a linear gradient of NaCl from 0 to 1 M in 20 column volumes. Fractions corresponding to the first peak were concentrated and stored in −80 °C.for further experiments.

### Crystallization

Crystallization was performed using the hanging-drop vapor diffusion method at 16 °C. For crystallization of Ups1/Mdm35 complex, seleno-methionine derived Ups1/Mdm35 complex or Ups1-Mdm35 fusion protein, 1 μl of the protein solution (20 mg/mL), 1 μl of the reservoir solution (8% tacsimate, pH5.0, 20%PEG3350), and 0.1 μl l of 2 M sodium thiocyanate were mixed and equilibrated against 200 μl of the reservoir solution. For crystallization of the Ups1/Mdm35-DHPA complex, purified Ups1/Mdm35 was incubated with 10-fold excess DHPA (830841C, Avanti Polar Lipids) in buffer C for 30 min at 4 °C. After incubation, the mixture was centrifuged at 17,000 g for 20 min at 4 °C. 1 μl of the supernatant (20 mg/mL) and 1 μl of the reservoir solution (30~35% tascimate, pH7.4, 0.1 M Bis-Tris propane, pH 7.5) were mixed and equilibrated against 200 μl of the reservoir solution.

### Data collection and structural determination

Diffraction data were collected at 100 K on beamline BL17U and BL18U of SSRF (Shanghai Synchrotron Radiation Facility) and processed with HKL2000 package ^28^. Single wavelength anomalous (SAD) diffraction data of the SeMet substituted Ups1/Mdm35 crystal was collected with a wavelength of 0.9791 Å. Selenium sites were determined using the program SHELXD ^29^. Phases were calculated and refined using SOLVE ^30^ and RESOLVE ^31^. An initial model was built using COOT ^32^ and further refined with PHENIX ^33^. Structures of Ups1/Mdm35-DHPA and Ups1-Mdm35 fusion protein were solved by the molecular replacement method and further refined with PHENIX. The quality of the structures was validated with MolProbity ^34^. The statistics of data collection and structure refinement are summarized in Supplementary Table 1.

### Liposome preparation

Phospholipids DOPC (1,2-dioleoyl-sn-glycero-3-phosphocholine, 850375C), DOPE (1,2-dioleoyl-sn-glycero-3-phosphoethanolamine, 850725C), DOPA (840875C) and 18:1-12:0 NBD-PA (1-oleoyl-2-(12-[(7-nitro-2-1,3-benzoxadiazol-4-yl)amino] dodecanoyl)-sn-glycero-3-phosphate, 810176C) were obtained from Avanti Polar Lipids. Rhodamine DHPE (L-1392) was purchased from Thermo Fisher Scientific. Lipids in stock solutions in chloroform were mixed at the desired molar ratio, and the solvent was evaporated. The lipid film was hydrated in an appropriate buffer. The lipid suspension was incubated at 37 °C for 30 min and then frozen in liquid nitrogen. This freeze-thaw cycle was repeated 5 times and then the thawed solution was extruded 21 times through a 0.1 μM filter using a mini-extruder (Avanti Polar Lipids).

### PA transfer activity assay

PA-transfer activities of the Ups1/Mdm35 complex and its mutants were measured by the fluorescent de-quenching assay as described previously ^8, 17^. Donor liposomes (6.25 uM; DOPC / DOPE / Rhodamine DHPE/18:1-12:0 NBD-PA = 50/40/2/8) were incubated with acceptor liposomes (25uM; DOPC / DOPE / DOPA = 50/40/10) in the presence or absence of 20 nM purified Ups1/Mdm35 complex and its mutants in 2 ml of assay buffer (20 mM Tris-HCl pH 7.5, 150 mM NaCl and 2 mM EDTA) at 25 °C. The NBD fluorescence was monitored using a fluorescence spectrometer (Hitachi F7000).

### Liposome co-sedimentation assay

Ups1/Mdm35 complex or its mutants were incubated with liposomes (DOPC: DOPA = 4:1) in the buffer (20 mM Tris-HCl pH 7.5, 150 mM NaCl and 1 mM EDTA) at a total volume of 50 μL at 25 °C for 30 min with the final concentration of 20 μM Ups1/Mdm35 and 8 mM liposomes, respectively. The mixture was then ultra-centrifuged at 250,000 g in a S140AT rotor (Doga Limited) on a micro ultracentrifuge (Hitachi Koki himac CS-FNX series) at 25 °C for 30min. Then the supernatants and pellets were subjected to 16.5% Tricine SDS-PAGE analysis. The stained gels were scanned and analyzed using Gel Doc™ EZ Gel Documentation System (Bio-rad).

### Molecular dynamics simulations and Data analysis

Multi-component lipid bilayers were constructed using CHARMM-GUI^35^ with a composition of DOPC/DOPE/DOPI/TOCL2/DOPA/DOPG/DOPS (8:6:2:1:1:1:1). Ups1_free_ and Ups1/Mdm35 complex systems were set up by placing the protein ∼8Å above the surface of the membrane. The systems were then solvated in explicit TIP3P ^36^ water molecules, with potassium and chloride ions in order to achieve a neutral and 0.18 M ionic solvent, using the solvate and auto-ionize tools of VMD ^37^. The MD simulations were performed using NAMD 2.11 package ^38^ and the CHARMM ^36^ force field^39, 40^ with CMAP correction^41^. Electrostatic interactions were calculated using the particle mesh Ewald sum method ^42^ with a cutoff of 12 Å. All hydrogen-containing covalent bonds were constrained by the SHAKE algorithm ^43^ except that SETTLE algorithm ^44^ was used for waters, therefore allowing an integration time step of 2 fs. Before production runs, the system was minimized in energy, heated to 310 K, and pre-equilibrated in the canonical ensemble while the protein backbone, and water oxygen atoms harmonically restrained with spring constant of 10 kcal mol^-1^ Å^2^. Simulations were then continued in the constant NPT ensemble (310K and 1 atm). Langevin thermostats with a damping coefficient of 0.5 ps^-1^ were used to control the system temperature. A Langevin-piston ^45^ barostat with a piston period of 2 ps and a damping time of 2 ps was used to control the pressure. Constraints were next released step-wise (with spring constant gradually decrease from 10 to 0 kcal mol^-1^ Å^2^) before starting the production runs. A total of 600 ns and 1040 ns of data were generated for Ups1/Mdm35 complex and Ups1_free_ systems respectively. Only data after reaching equilibrium would be taken for further analysis.

## DATA AVAILABILITY

The coordinates and structural factors of crystal structures of SeMet substituted UPS1/Mdm35, the fusion form and the DLPA bound state have been deposited into Protein Data Bank with the accession codes 5JQL, 5JQM and 5JQO, respectively.

## AUTHOR CONTRIBTUIONS

F. S. and Y.Z. initiated and supervised the project. J.L. performed molecular cloning, protein expression, purification, crystallization, data collection and structure determination. K.C. and J.F. performed molecular dynamics simulations. L.Y. performed mutagenesis, biophysical assays and PA transfer activity assay. Y.Z. analyzed the data. Y.Z., J.F. and F.S. wrote the manuscript.

## ACKNOWLEDGMENTS

We would like to thank Ping Shan and Ruigang Su (F.S. lab) for their assistances of lab management. We are grateful to Qiangjun Zhou (F.S. lab) for his help in structure determination. We would like also to thank the engineers from Core Facility of Protein Sciences, Institute of Biophysics, for their help in ultra-centrifuge and fluorescence spectrometry experiments. This work was supported by grants from Ministry of Science and Technology of China (2017YFA0504700) to F.S. and Natural Science Foundation of China (31771566) to Y.Z.

## Supplementary Figures

**Supplementary Figure 1.**
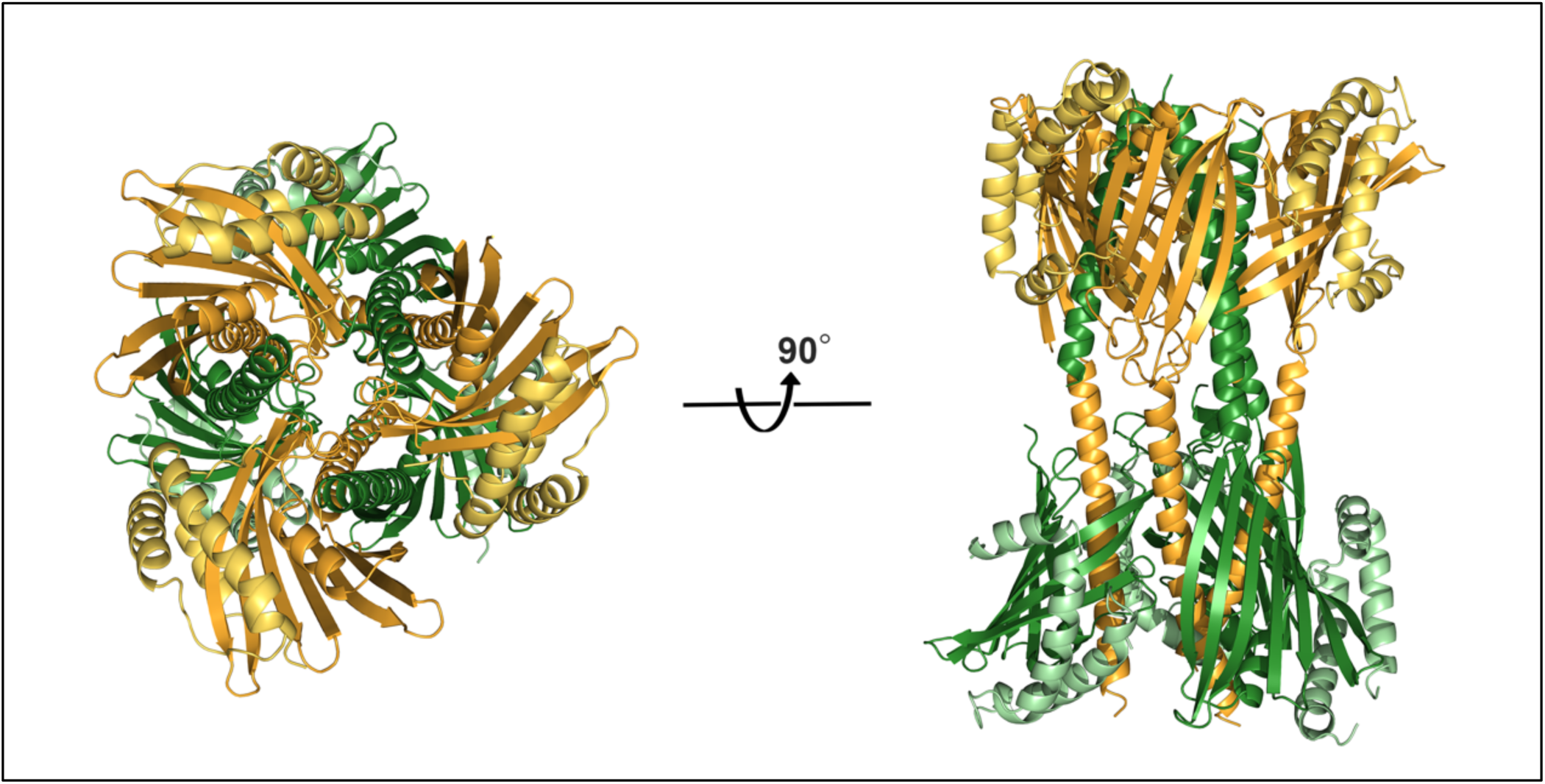
The crystallographic asymmetry unit of SeMet-substituted Ups1/Mdm35 crystal structure (PDB code 5JQL) that contains six Ups1/Mdm35 heterodimers.

**Supplementary Figure 2.**
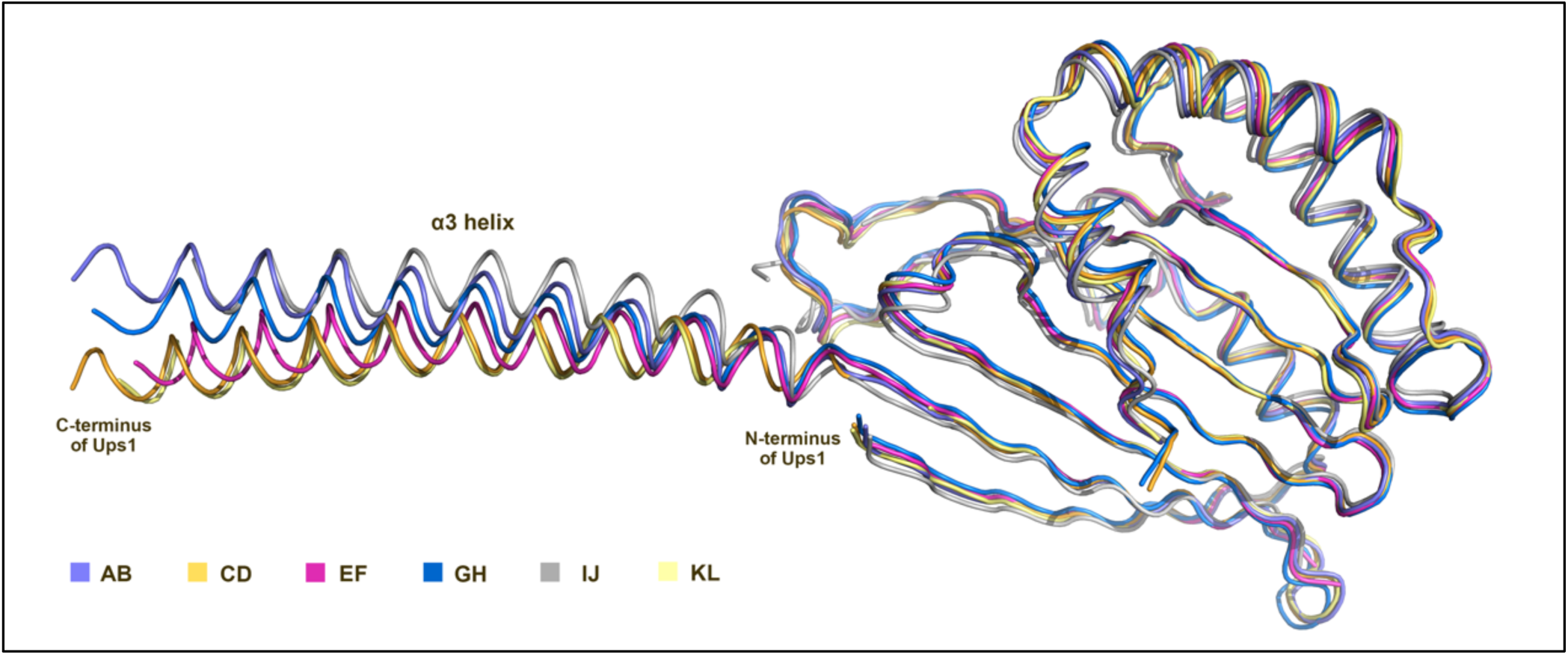
Superposition of six Ups1/Mdm35 molecules in an asymmetric unit of SeMet-substituted Ups1/Mdm35 crystal structure (PDB code 5JQL). AB, CD, EF, GH, IJ and KL represent six Ups1/Mdm35 molecules respectively.

**Supplementary Figure 3.**
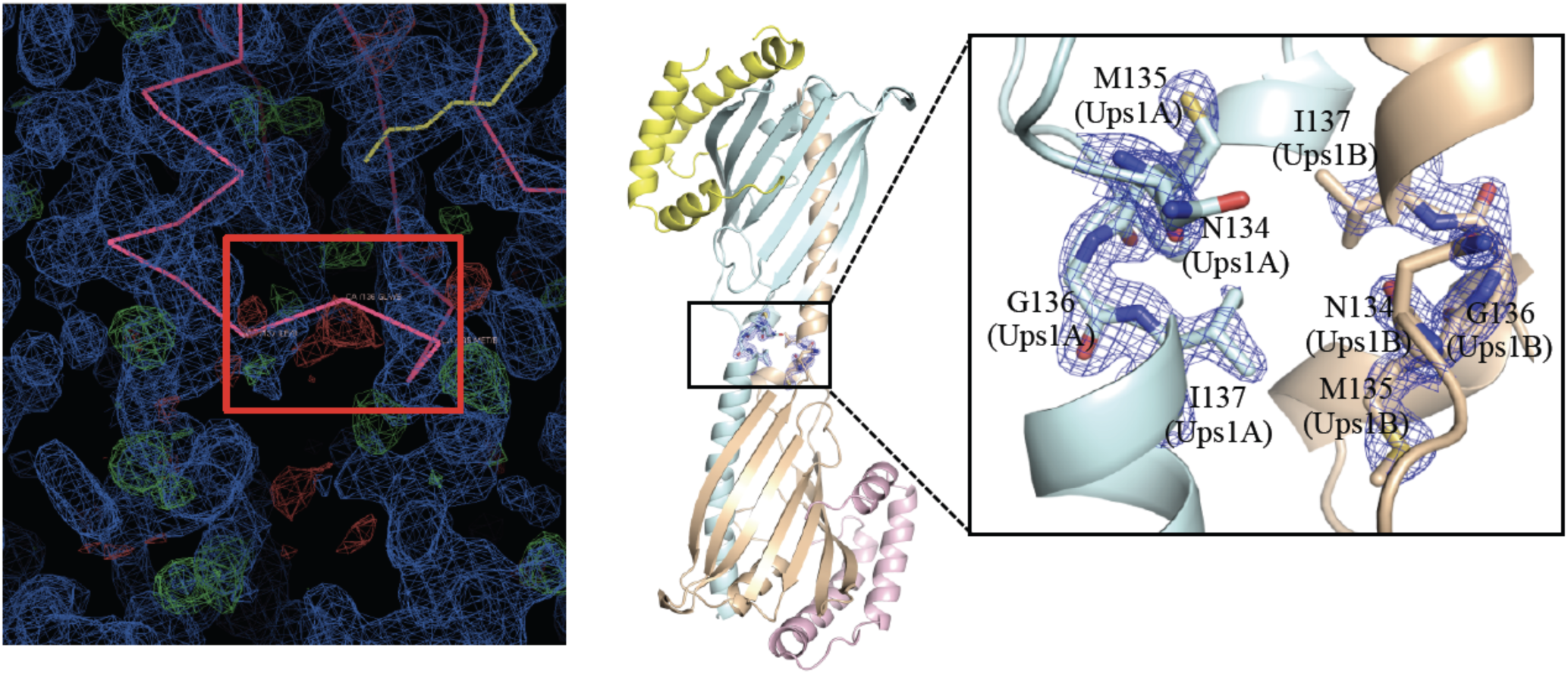
Rebuilding of residues N134-I137 of Ups1 in the Ups1/Mdm35-DLPA structure (PDB: 4XIZ). **Left:** The electron density map and the Cα backbones around residues N134-I137 of Ups1. Ups1/Mdm35 is colored in pink and DLPA is colored in yellow. 2FoFc map is colored in blue and FoFc difference map is colored in green and red. The Cα backbones of N134-I137 that do not fit into the electron density are highlighted by a red box. **Right:** Rebuilt structure of Ups1/Mdm35-DLPA. Ups1 molecules are colored in light blue and wheat. Mdm35 molecules are colored in yellow and pink. The magnified view shows the rebuilt Cα backbones of N134-I137, which fit well into the electron density (2FoFc map contoured at 2.0 σ and colored blue).

**Supplementary Figure 4.**
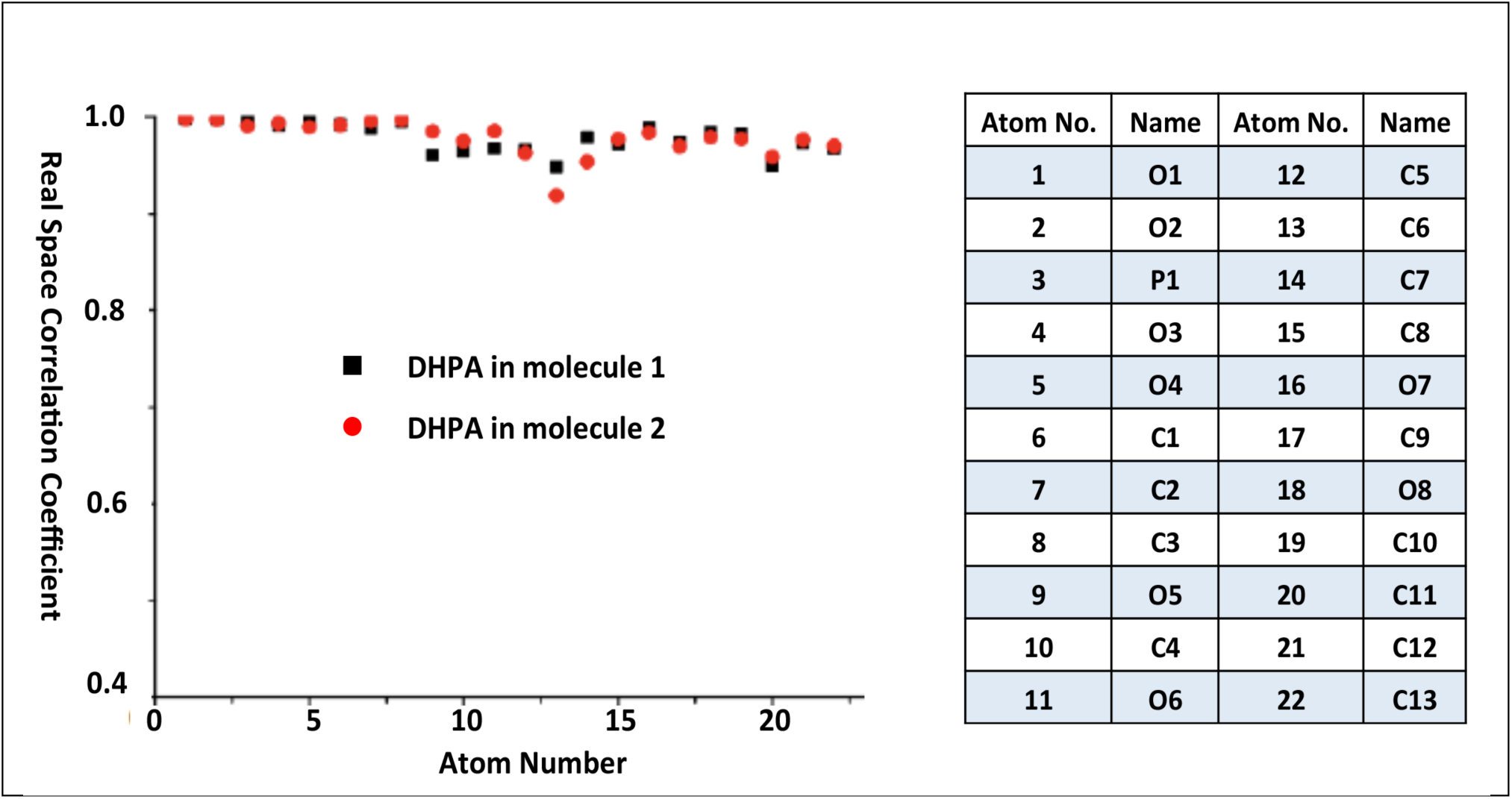
Real-space correlation coefficients (RSCC) as a function of atom of two DHPA molecules in an asymmetric unit in the Ups1/Mdm35-DHPA structure (PDB code 5JQO). No atom has a coefficient below 0.9 indicating that the ligands fit well into the electron density.

**Supplementary Figure 5.**
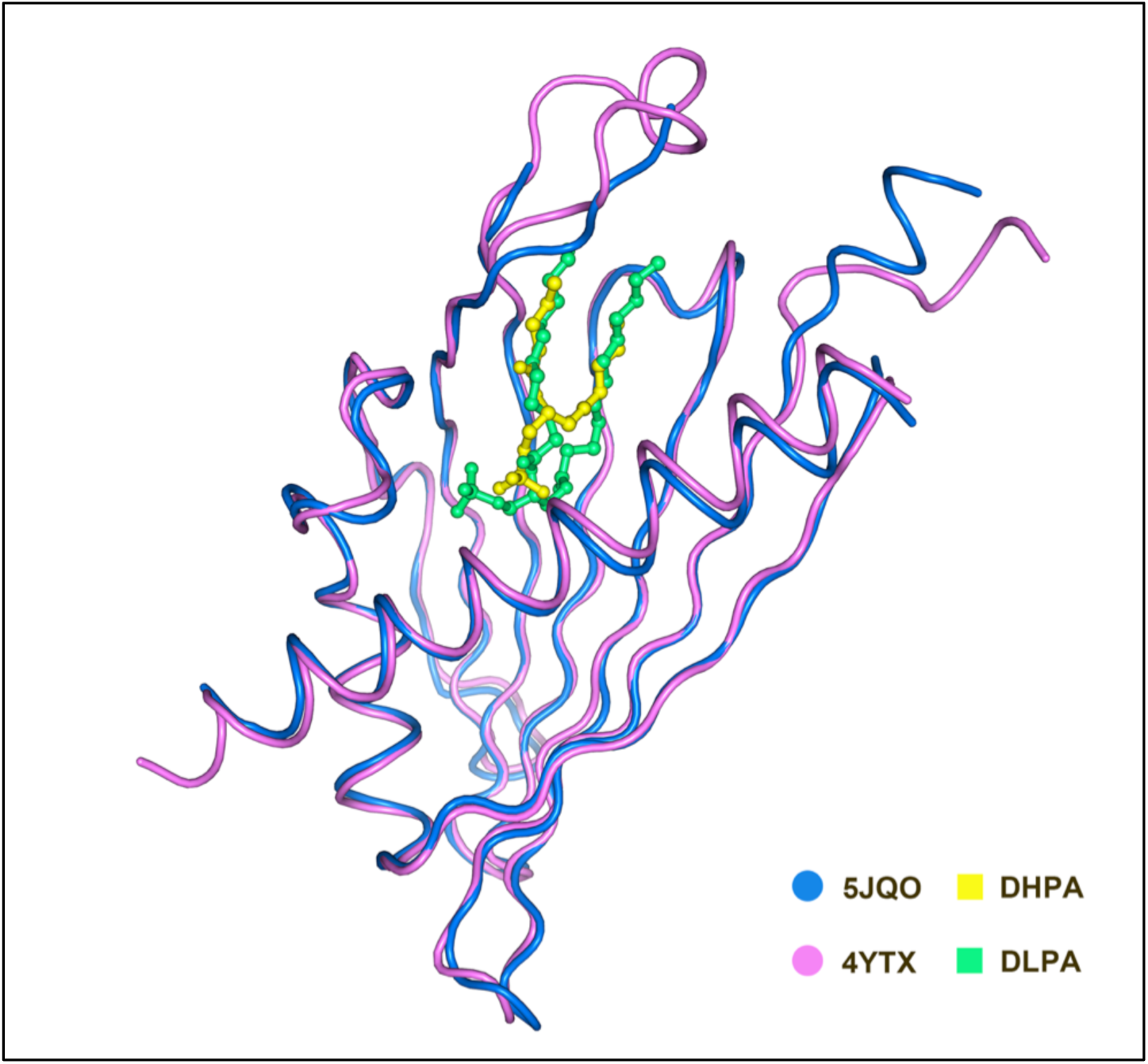
Structural comparison of Ups1/Mdm35-DHPA (PDB code 5JQO) and Ups1/Mdm35-DLPA (PDB code 4YTX).

**Supplementary Figure 6.**
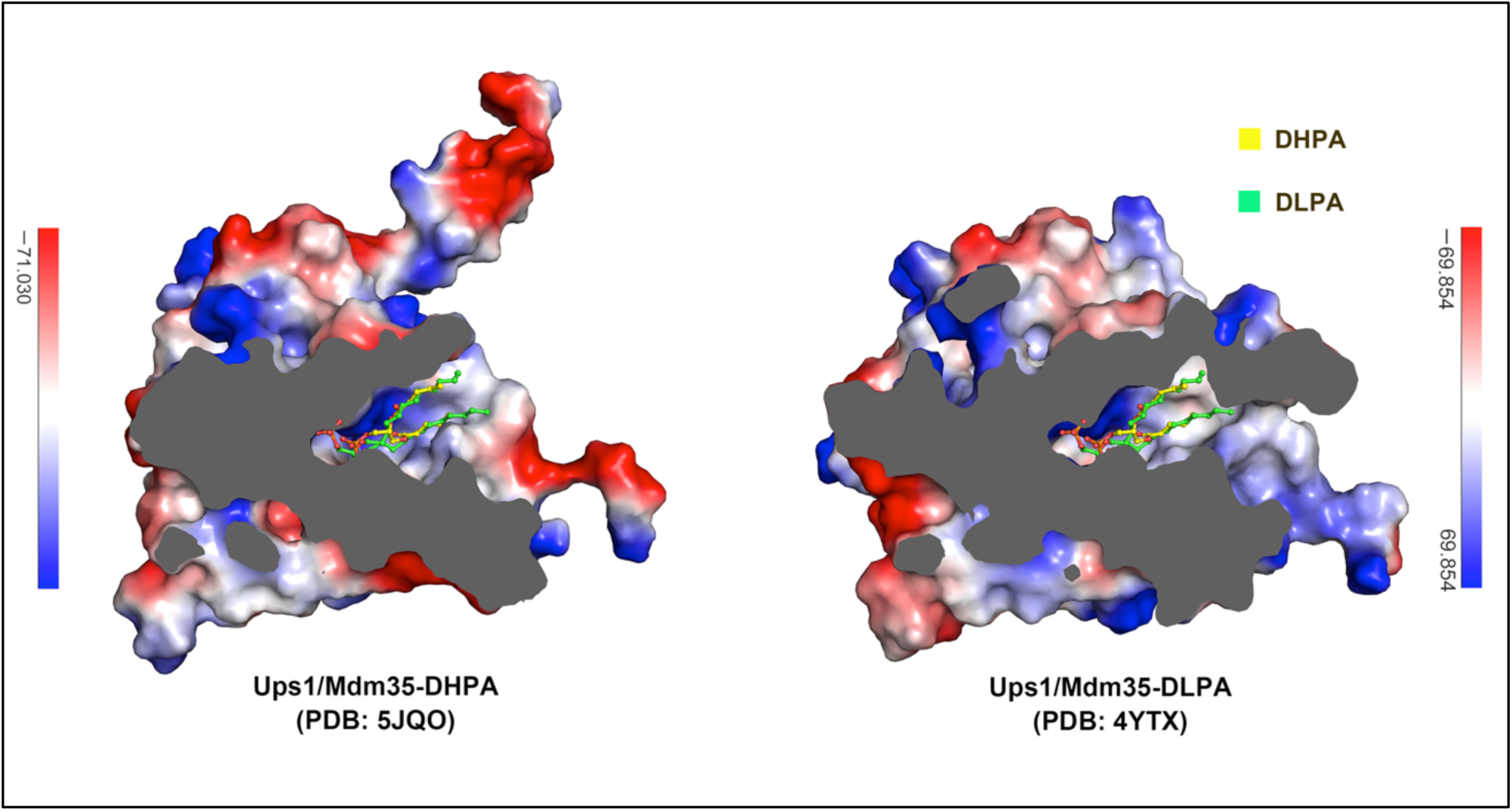
Comparison of binding sites of DHPA and DLPA in Ups1/Mdm35 structures. Surface electrostatic potential maps of Ups1/Mdm35 structures are cutaway to show the DHPA/DLPA binding pocket. DHPA and DLPA molecules are shown as stick models.

**Supplementary Figure 7.**
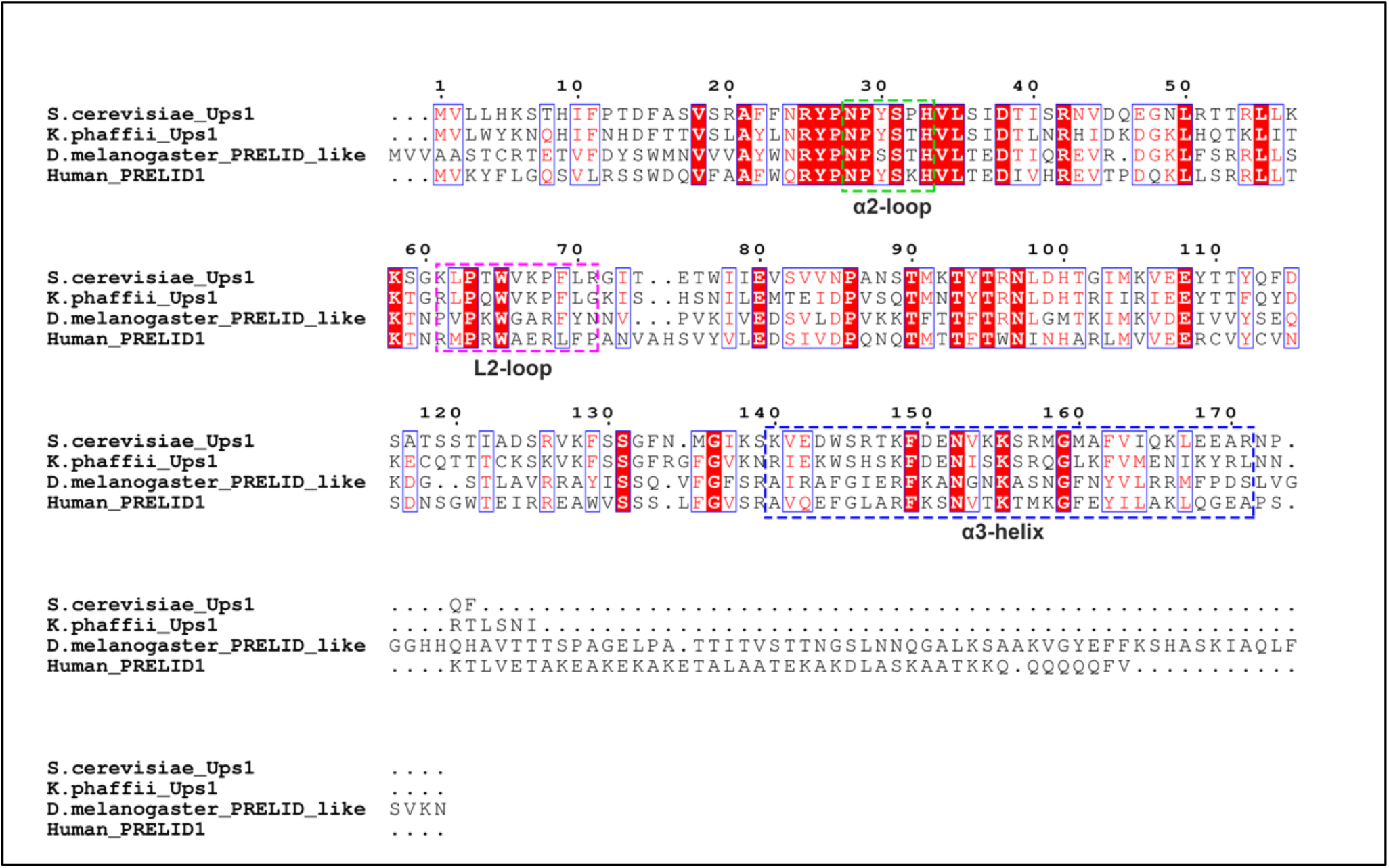
Sequence alignment of Ups1 homologues. Invariant residues are shaded in red, conserved residues are colored in red and framed in blue box. The membrane-binding elements of Ups1, including the α2-loop, L2-loop and α3-helix, are framed as indicated.

**Supplementary Figure 8.**
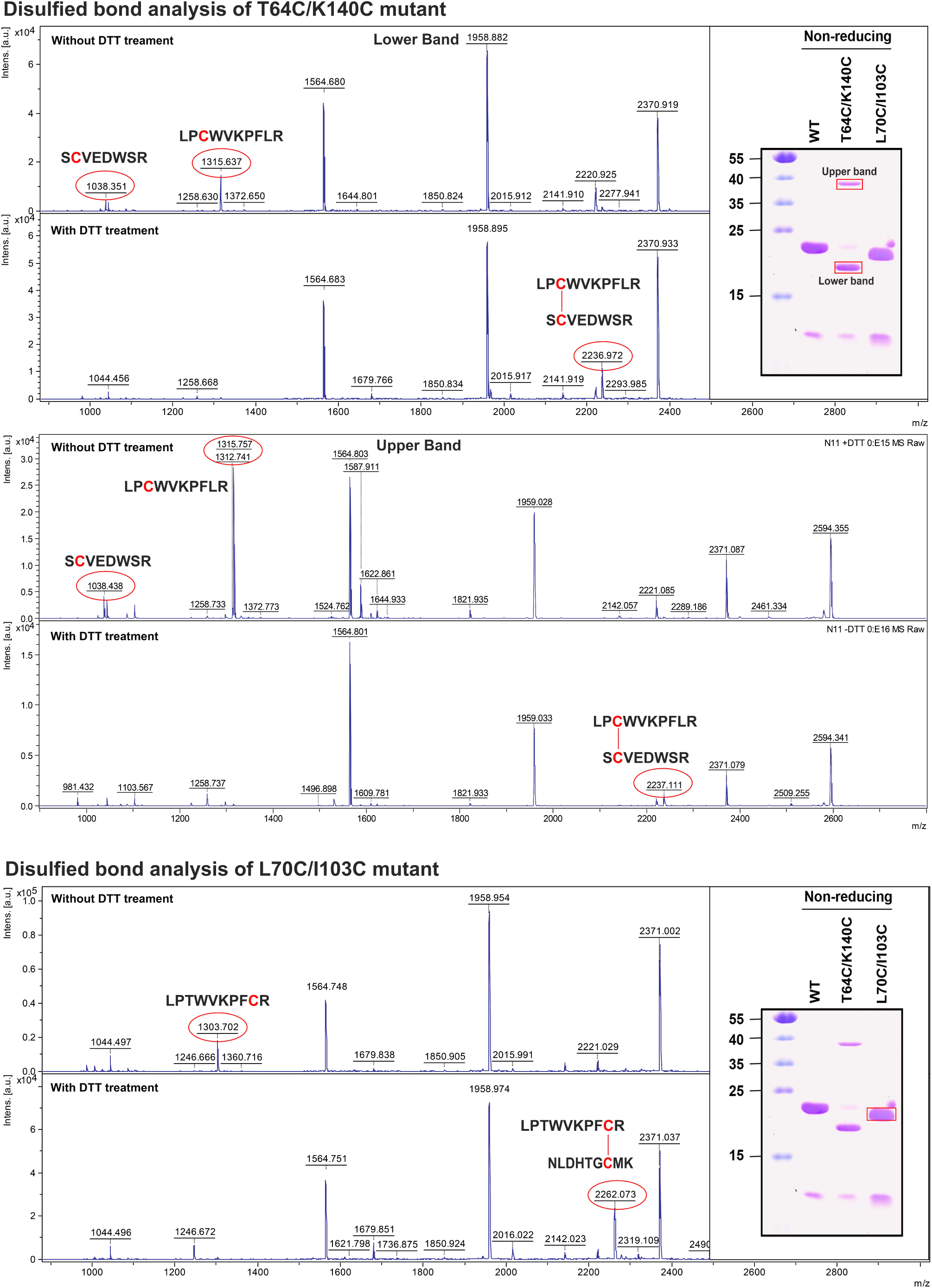
Mass spectrometry analysis of the disulfide bonds in T64C/K140C and L70C/I103C mutants. The MALDI-TOF (matrix assisted laser desorption ionization time of fly) mass spectra of the target bands are listed as indicated.

**Supplementary Figure 9.**
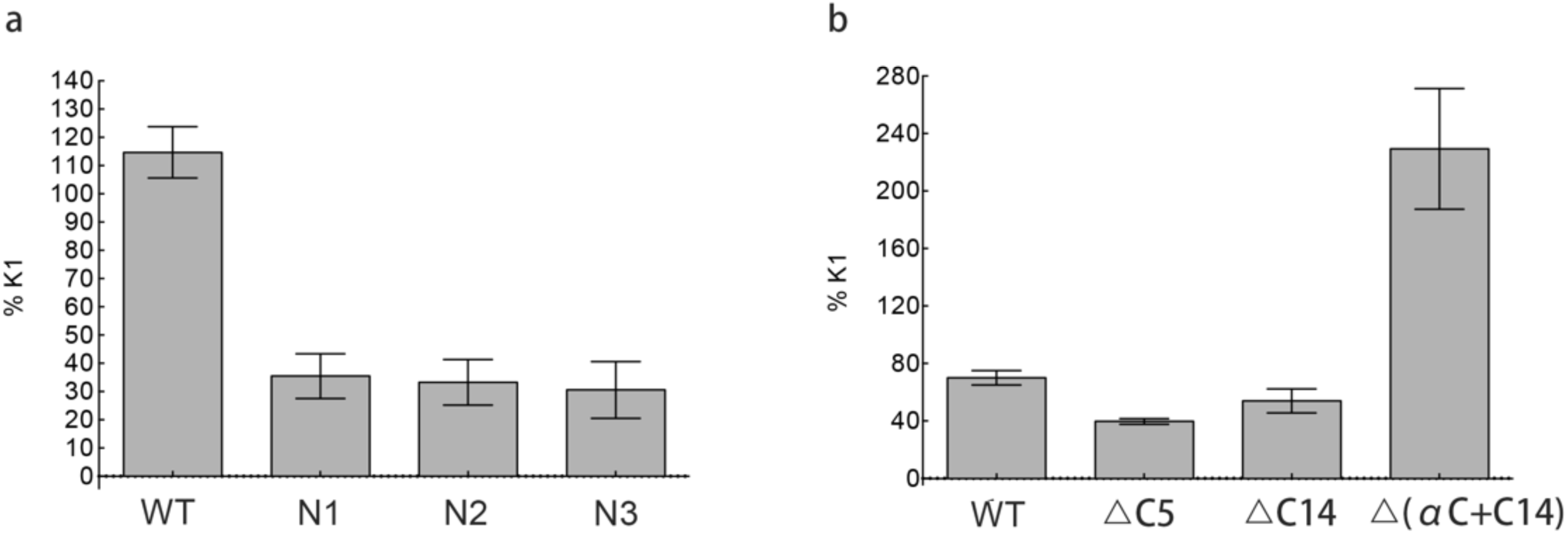
K1 values of the indicated mutants (see Figs. 2 and 3). K1 is calculated according to the equation (25) of Supplementary Text. N1, the mutant of Ups1-2E; N2, the mutant of Ups1-3E; N3, the mutant of Ups1-4E.

## Supplementary Text

**Kinetic equations of Ups1/Mdm35-mediated PA transport and the derivation of K1 constant from the liposome co-sedimentation assay.**

When the reactions (a) and (b) reach equilibrium, ten equations can be derived as follows,

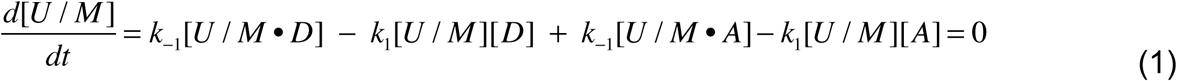

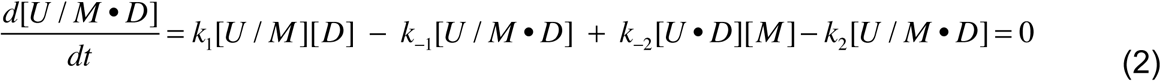

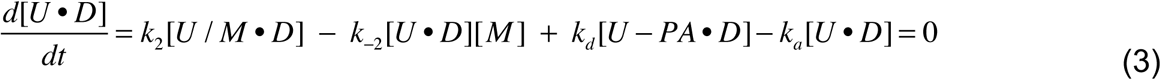

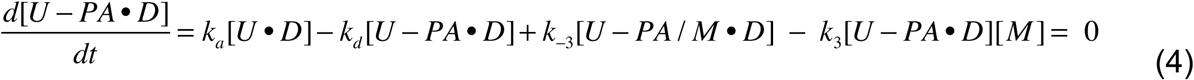

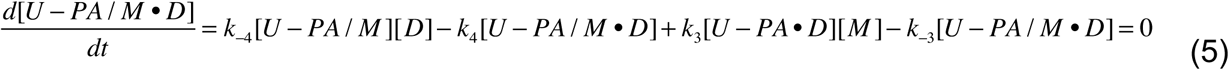

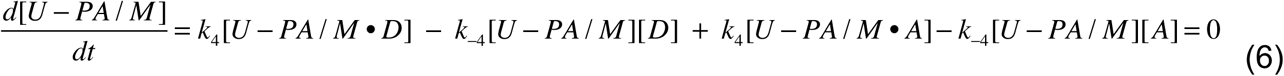

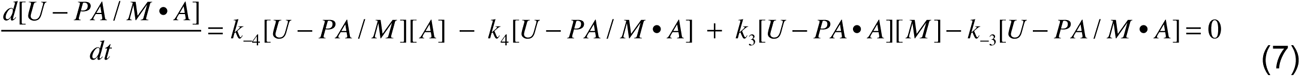

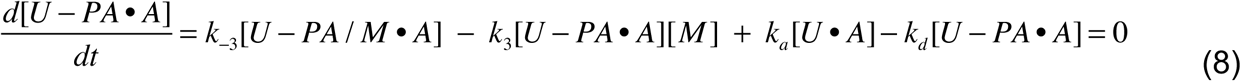

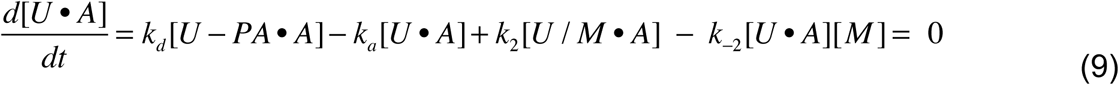

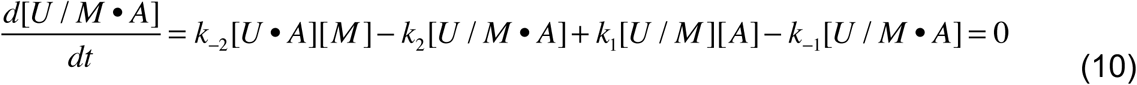

In liposome co-sedimentation assay, donor and acceptor membrane are identical, (1) can be simplified as

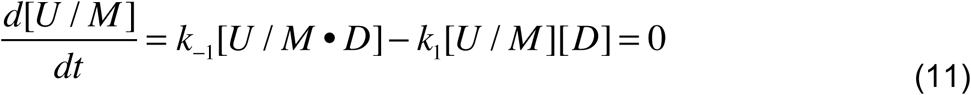

Substitution of (11) into (2), (3), (4) and (5) yields the following equation:

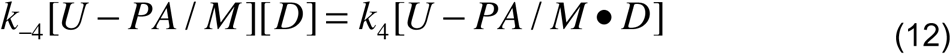

We assume k_1_ ≈ k_-4_ and k_-1_ ≈ k_4_, (12) can be rewritten into:

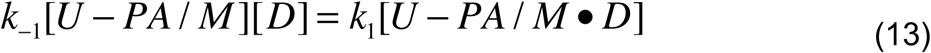

Addition of (11) and (13) yields,

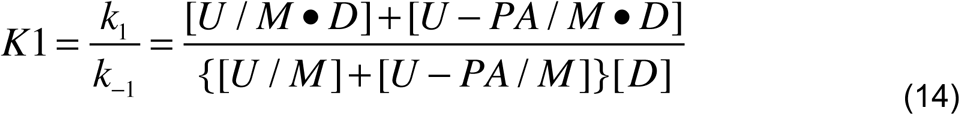

In liposome co-sedimentation assay, we have,

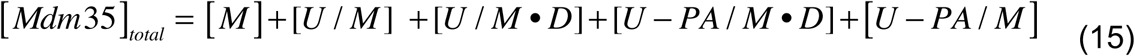

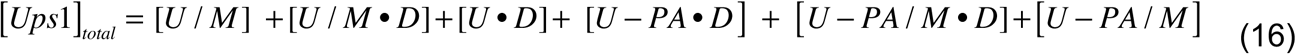

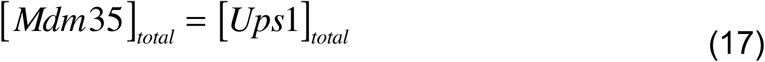

Based on the scheme of Figure 2g, the following equations can be deduced:

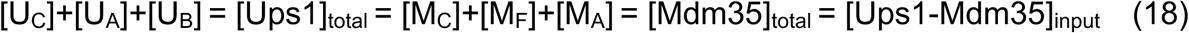

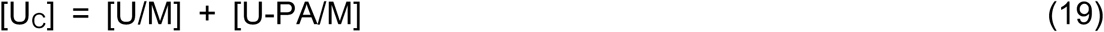

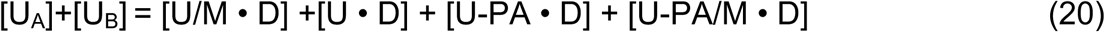

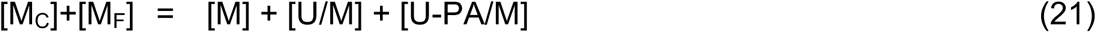

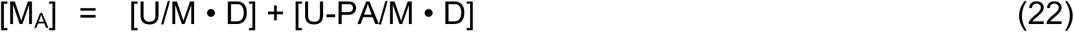

Substitution of (19) and (22) into (14) yields the following equation,

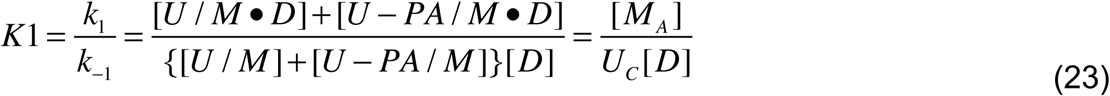

---

**Supplementary Table 1.** Statistics of data collection and structure refinement.

|  | SeMet labelled<br>Ups1/Mdm35 | Ups1-Mdm35 | Ups1/Mdm35-DHPA |
| --- | --- | --- | --- |
| Data collection |  |  |  |
| Space group | $P2_1$ | $C2$ | $P4_1$ |
| Cell dimensions |  |  |  |
| a,b,c (Å) | 83.3,130.7,84.3 | 118.6,68.3,105.1 | 90.1,90.1,81.3 |
| $\alpha,\beta,\gamma$ (°) | 90,103.1,90 | 90,102.1,90 | 90,90,90 |
| Data set |  |  |  |
| Wavelength (Å) | 0.9791 | 0.9791 | 0.9786 |
| Resolution range (Å) | 50 ~ 2.90 | 50 ~ 1.50 | 50 ~ 3.55 |
| Outer Shell (Å) | 2.95 ~ 2.90 | 1.53 ~ 1.50 | 3.61 ~ 3.55 |
| $R_{\text{merge}}$ | 0.074(0.438) | 0.084(0.42) | 0.053(0.67) |
| $R_{\text{pim}}^a$ | 0.045(0.266) | 0.052(0.279) | 0.022(0.273) |
| $I/\sigma I$ | 31.7(4.2) | 37.7(3.0) | 37.8 |
| Completeness (%) | 98.4(97.0) | 98.6(88.9) | 100(100) |
| Redundancy | 3.6(3.6) | 3.6(3.1) | 6.7(6.9) |
| CC1/2 high<br>resolution shell | 0.903 | 0.820 | 0.849 |
| Refinement |  |  |  |
| No.reflections | 39282 | 129414 | 7762 |
| $R_{\text{work}}/R_{\text{free}}^b$ (%) | 20.0/24.8 | 16.4/20.2 | 28.6/32.7 |
| No of atoms |  |  |  |
| protein | 11018 | 5610 | 3281 |
| Ligand | — | — | 44 |
| water | — | 811 | — |
| $B$-factors (Å<sup>2</sup>)<sup>c</sup> | | | |
| Protein | 38.1 | 23.0 | 63.3 |
| Ligand | — | — | 82.8 |
| water | — | 35.1 | — |
| r.m.s.d.<sup>d</sup> |  |  |  |
| Bond lengths (Å) | 0.01 | 0.007 | 0.013 |
| Bond angles (°) | 1.3 | 1.1 | 1.78 |
| Ramachandran |  |  |  |
| Favored (%) | 98.2 | 99.3 | 99.3 |
| Allowed (%) | 1.8 | 0.7 | 0.5 |
| PDB ID | 5JQL | 5JQM | 5JQO |

Values in parentheses refer to the highest resolution shell. —, not available. <sup>a</sup> $R_{\text{pim}}$ is the precision-indicating merging R factor. It measures the quality of the data after averaging the multiple measurements. <sup>b</sup> $R_{\text{work}} = \sum(|F_{\text{p}}(\text{obs})| - |F_{\text{p}}(\text{calc})|) / \sum|F_{\text{p}}(\text{obs})|$ ; $R_{\text{free}}$ = R factor for a selected subset (5%) of the reflections that was not included in prior refinement calculations. <sup>c</sup> Averaged B-factors of corresponding atoms in both chains from one asymmetric unit are shown separately. <sup>d</sup> Root mean square deviation from ideal values.

## Supplementary Movies

**Supplementary Movie 1. Molecular dynamics simulation of Ups1_free_ on the lipid bilayer.** Secondary structural elements are shown in cartoon. Alpha helices are colored in magenta and beta strands in yellow.

**Supplementary Movie 2. Molecular dynamics simulation of Ups1/Mdm35 complex on the lipid bilayer.** Secondary structural elements are shown in cartoon. Ups1 is colored in cyan and Mdm35 in gold.

**Supplementary Movie 3. Molecular dynamics simulation of Ups1_free_ on the lipid bilayer from 800 ns to 1023 ns.** A front view of the membrane binding interface of Ups1 is presented here.

**Supplementary Movie 4.** Apo-Ups1/Mdm35 can adopt at least two conformations (4YTW and 5JQM) in solution and switch freely between them.In 5JQM, the interaction between W65 and the hydrophobic patch composed of I137, V141 and W144, is stronger than in 4YTW. When Apo-Ups1/Mdm35 approaches the lipid bilayer, Ups1/Mdm35 intends to interact with the membrane through the membrane-binding residues of Ups1 as a whole. Then a thermodynamic perturbation of the system will induce the dissociation of Mdm35 from Ups1, which begins with the αC-helix detachment. The resulting allosteric effects alleviate the flexibility of the membrane-binding residues of Ups1 and leads to the insertion of F69 into the membrane (MD simulated Ups1,350ns). At 1023ns, the head of DLPA interacts with Ups1 (MD simulated Ups1,1023ns). Then DLPA is trying to enter the PA-binding pocket of Ups1 (X-state Ups1, 4YTX-KJ). Finally, DLPA enters the pocket completely (DLPA-bound Ups1, 4YTX-AB).

